# A mitochondrial quality control mechanism reverses the phagosome maturation arrest caused by *Mycobacterium tuberculosis*

**DOI:** 10.1101/2023.12.01.569475

**Authors:** Surbhi Verma, Shweta Thakur, Mrinmoy Das, Raman D. Sharma, Vikas Yadav, Priya Sharma, Mardiana Marzuki, Shihui Foo, Giulia M. Piperno, Mehak Z. Khan, Babu Mathew, Meenu Bajpai, Jaswinder Singh Maras, Shanshan Howland, Sovan Sarkar, Federica Benvenuti, Amit Singh, Vinay Nandicoori, Amit Singhal, Dhiraj Kumar

## Abstract

Phagosome maturation arrest (PMA) imposed by *Mycobacterium tuberculosis* (*Mtb*) is a classic tool that helps *Mtb* evade macrophage anti-bacterial responses. The exclusion of RAB7, a small GTPase, from *Mtb*-phagosomes causes PMA. Here, we report an unexpected mechanism that triggers crosstalk between the mitochondrial quality control (MQC) and the phagosome maturation pathways that reverses the PMA. CRISPR-mediated p62/SQSTM1 depletion (*p62^KD^*) does not appear to impact mitochondrial quality. The *p62^KD^* cells are restrictive to *Mtb* growth, triggered by an increasingly oxidative environment and increased lysosomal targeting. The lysosomal targeting of *Mtb* is facilitated by enhanced TOM20^+^ mitochondria-derived vesicles (MDVs) biogenesis, a key MQC mechanism. In *p62^KD^* cells, TOM20^+^-MDVs biogenesis is MIRO1/MIRO2-dependent and gets delivered to lysosomes for degradation in a RAB7-dependent manner. Upon infection in *p62^KD^* cells, TOM20^+^-MDVs get extensively targeted to *Mtb*-phagosomes, inadvertently facilitating RAB7 recruitment, PMA reversal and lysosomal targeting of *Mtb*; the phenotype also replicated in p62/SQSTM1 knockout cells. Triggering MQC collapse in *p62^KD^*cells further diminishes *Mtb* survival, signifying cooperation between redox- and lysosome-mediated mechanisms. The MQC-anti-bacterial pathway crosstalk could be exploited for host-directed anti-tuberculosis therapies.

## Introduction

Tuberculosis (TB), caused by the pathogen *Mycobacterium tuberculosis* (*Mtb*), continues to pose challenges to global communicable disease management and is one of the leading causes of mortality due to infectious diseases. The ability of the bacteria to alter and/or hijack various host cellular responses in its favour is one of the major factors contributing to the success of *Mtb* as a human pathogen. Arguably, understanding the host physiological processes hijacked or altered by *Mtb* also presents an opportunity to develop unconventional strategies to target and control the infection. In the past decade, several host-dependency factors have been identified, through large genome-wide screens, high-throughput transcriptome and systematic analysis of bacterial trajectory inside the host macrophages, the primary cells infected by *Mtb* (1).

The role of lysosomes in killing bacterial pathogens, including *Mtb* is well-known (2). Macrophages phagocytose bacterial pathogens, forming phagosomes, which mature from early to late phagosomes, eventually fusing with the lysosomes, where the pathogen is degraded (3). The phagosome maturation involves RAB5 (early phagosome) to RAB7 (late phagosome) exchange as a critical rate-limiting step (4). *Mtb* is known to block this RAB-conversion by excluding RAB7 recruitment, which helps them evade the lysosomal killing (5). Several mechanisms, including the role of lipid mediators and other signalling events, have been shown to contribute towards the phagosome maturation arrest (PMA) caused by *Mtb* (6).

Autophagy is another important anti-bacterial process that kills bacteria through a lysosomal targeting mechanism (7, 8). Autophagy performs crucial housekeeping functions like degradation of misfolded proteins or damaged organelles like mitochondria, peroxisomes, Endoplasmic Reticulum etc., in addition to its role in response to stresses like oxidative, nutritional or infection-induced stress (9). Understanding how autophagy machinery distinguishes and segregates its homeostatic cargos, as mentioned above, from those like pathogens is of considerable fundamental significance (10).

The cargos targeted for autophagic degradation are usually ubiquitinated, followed by their recognition by the autophagy adaptors, which interact with the ubiquitinated cargos and ATG8 or LC3B to capture the cargos into the growing autophagosomes (11). Autophagy adaptors are a family of proteins with ubiquitin-binding domains and LC3/ATG8 interaction domains. These adaptors primarily include p62/SQSTM1, NDP52, NBR1, OPTN and TAX1BP1 (12). Each of these adaptors is reported to have its roles defined in cargo selectivity. However, some reports suggest functional redundancy among them (13, 14).

While autophagic targeting of bacteria via xenophagy is part of the defence arm of autophagy, degradation of cellular cargos like mitochondria via mitophagy constitutes the homeostatic arm (10, 15). Mitophagy is a key pathway for mitochondrial quality control (MQC) through which damaged mitochondria are removed from the cell (16). Since mitochondrial damage is a normal consequence of respiration and other oxidative stresses, their quality control through mitophagy and other pathways is fundamental to cellular survival (17, 18).

In this study, while studying the effect of depletion of autophagy adaptor proteins like p62/SQSTM1 on intracellular *Mtb* survival, we unravel an unexpected MQC that facilitated *Mtb* targeting to the lysosomes by reversing the PMA. Our data provide mechanistic insights on the crosstalk between MQC and conventional host anti-bacterial pathways that are of both fundamental and biomedical relevance.

## Results

### Selective recruitment of autophagy adaptors to *Mtb* phagosomes and mitochondria

Since the autophagy adaptors target the autophagic cargo during selective autophagy, we investigated how the expression and recruitment of these adaptors to the phagosomes and mitochondria are affected upon *Mtb* infection. We first analysed for any perturbation in the expression of the major autophagy adaptors, such as *p62/SQSTM1*, *OPTN*, *NBR1*, *NDP52* and *TAX1BP1*, in THP-1 macrophages upon infection with the *Mtb* strain, H37Rv (MOI 1:10, see methods). For all the infection experiments in this study, the time 0 hpi (hours post-infection) includes incubation with H37Rv for 4 hours followed by 2 hours of amikacin treatment to kill extracellular bacteria (detailed in the methods section). At the transcript level, *OPTN* and *p62/SQSTM1* showed increased expression at 0-, 24- and 48-hpi (Fig. S1A). However, immunoblotting indicated higher expression of only p62/SQSTM1 proteins upon infection (Fig. S1B-C). To target the bacteria for autophagic degradation, autophagy adaptors need to get recruited onto the bacteria or bacterial phagosomes. Therefore, we evaluated the recruitment of autophagy adaptors on mCherry-H37Rv phagosomes by immunostaining and confocal microscopy at 0-, 6-, 12-, 24- and 48-hpi (see methods). While all of the five autophagy adaptors being studied were recruited onto the bacterial phagosomes (Fig.1A-E), we did observe some selectivity in the extent of colocalisation. For instance, OPTN showed the least recruitment to bacterial phagosomes, ranging from 2-4% colocalisation across all the time points (Fig. 1B), whereas p62/SQSTM1 showed maximum recruitment starting at ∼12% colocalisation at 0 hpi and increasing to ∼40% colocalisation at 48-hpi (Fig. 1A). The other three adaptors showed intermediate levels of recruitment: NBR1 5-20%, NDP52 10-15% and TAX1BP1 5-15% (Fig. 1C-E). Notably, only p62/SQSTM1 and NBR1 showed increased recruitment to the bacterial phagosomes over time as the infection progressed (Fig. 1A, 1C).

**Fig. 1.**
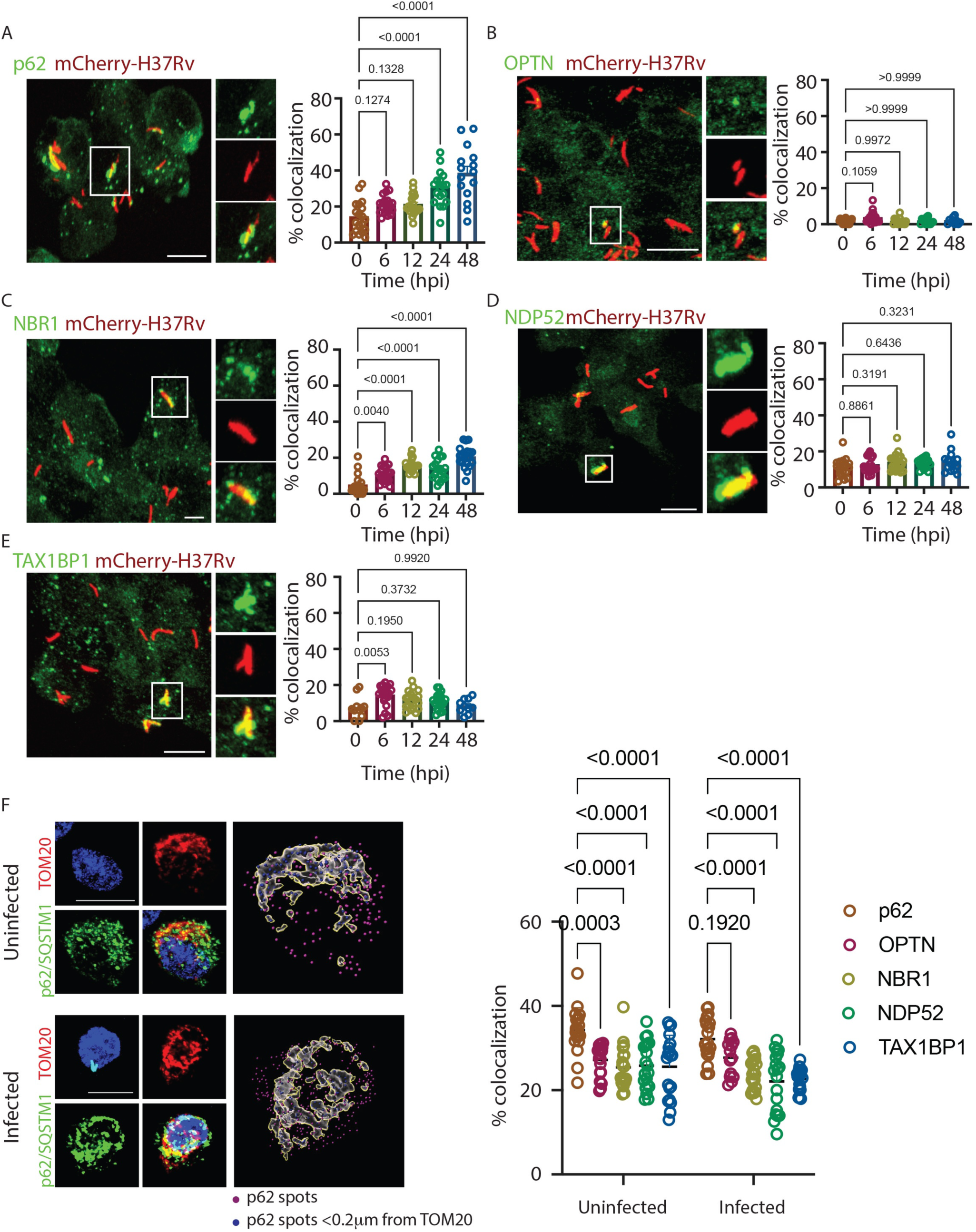
Selective recruitment of autophagy adaptors to *Mtb* phagosomes and mitochondria: **(A-E)** Confocal images show the colocalisation of individually immunostained autophagy adaptors as indicated (green) with mCherry-H37Rv (red) respectively at indicated time points post-infection in THP-1 macrophages. The area in the box is zoomed in and shown across the individual colour channels at the right. The bar graphs at the right represent the percent co-localization calculated manually using Imaris software. Scale bar: 10 μm. Data show mean ± SEM, *n* >10 fields per group, 20-35 cells per field, from three independent experiments. **(F)** Representative images of the interaction between immunostained p62/SQSTM1 (green) and mitochondrial marker TOM20 (Red). Nucleus were stained with DAPI (blue) at 24 hpi in THP-1 macrophages. To the right, representative 3D reconstruction is shown for TOM20 structures and p62/SQSTM1 spots. Magenta spots: non-interacting, >0.2 μm from TOM20, blue spots: interacting, <0.2 μm from the TOM20 structure. The graph represents the percentage of indicated adaptor spots interacting with TOM20 structures, calculated by the shortest distance module of Imaris software. Scale bar: 10 μm. Data show mean ± SD, n=15 fields per group, 20-35 cells per field, from three independent experiments. Scale Bar: 10 μm.

Since there are common autophagy adaptors involved in the autophagic targeting of bacteria (for xenophagy) as well as other cellular cargos like mitochondria (for mitophagy) (10, 19), we studied the distribution of these autophagy adaptors to a major cellular autophagic cargo such as mitochondria, during infection. For this, we analysed TOM20, an integral outer membrane mitochondrial protein that can be reliably used to track mitochondrial dynamics inside the cell (20). In uninfected as well as mCherry-H37Rv infected THP-1 cells, the recruitment of p62/SQSTM1 to TOM20^+^ mitochondria was significantly higher than any other autophagy adaptors (Fig. 1F). Upon H37Rv-infection, the pattern of autophagy adaptor recruitment on TOM20^+^ mitochondria mainly remained unchanged, except that there was a marginal increase in OPTN recruitment. Thus, no significant re-distribution of autophagy adaptors on mitochondria occurred during *Mtb* infection.

### Role of autophagy adaptors in intracellular bacterial growth and survival

All five autophagy adaptors being studied were recruited to the bacteria and mitochondria in the macrophages, albeit differentially. To investigate their involvement in the bacterial targeting for autophagic degradation, we individually depleted the autophagy adaptors in the H37Rv-infected macrophages. Each one of the five autophagy adaptors (*p62/SQSTM1, OPTN, NBR1, NDP52* and *TAX1BP1*) was knocked down in separate experiments using siRNAs (see methods) with 60-80% knockdown efficiency (Fig. S2A). While different autophagy adaptors showed differential recruitment to the phagosomes (Fig. 1), surprisingly, all except *NDP52* knockdown resulted in a lower intracellular bacterial CFU (Fig. 2A).

**Fig. 2.**
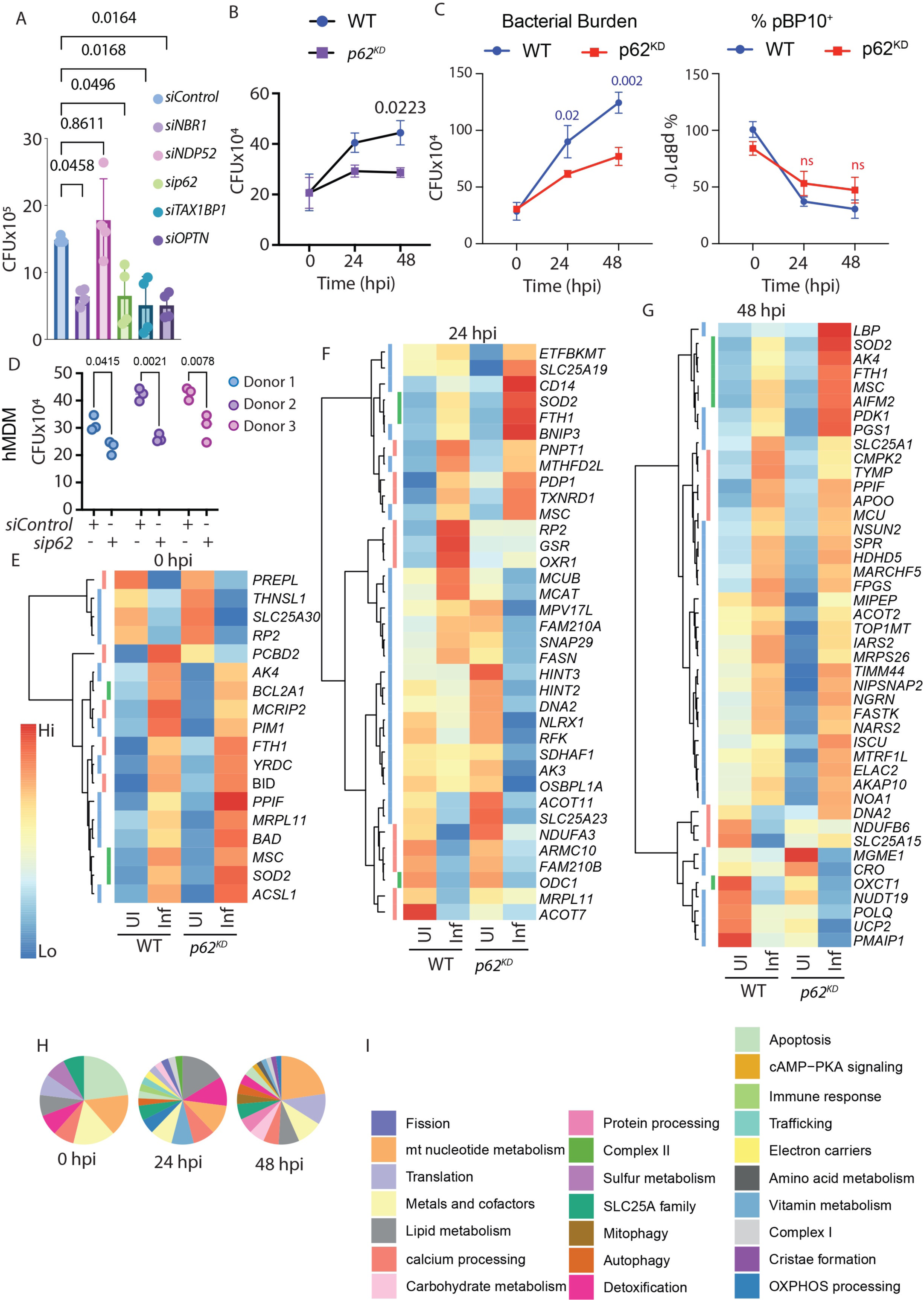
Autophagy adaptors regulate intracellular *Mtb* survival: **(A)** The bar graph represents the H37Rv CFU upon individual autophagy adaptor depletion using siRNAs in THP-1 macrophages 24 hpi. Data show mean ± SD, from four independent experiments. **(B)** Time course assay of H73Rv CFU in WT and *p62^KD^* THP-1 macrophages. Data show mean ± SD from four independent experiments. **(C)** WT and *p62^KD^* cells were infected with pBP10^+^-H37Rv, and CFU plating was done at designated time points on kan^+^ or Kan^-^ 7H11 plates. The total bacterial burden (i.e., CFU on Kan^-^ plates, left panel) and %pBP10^+^*Mtb* (ratio of CFU on Kan^+^ to Kan^-^ plates, right panel) for the WT (left) and *p62^KD^* (right) cells are shown. Data is mean± SD, representative from two independent experiments. **(D)** The bar graph represents H37Rv CFU in hMDMs in control and p62 knockdown conditions at 24 hpi. Data show mean ± SD; individual data points for three different donors are shown. **(E-G)** Heatmap represents the mean z-scaled expression of mitochondrial-related DEGs (differentially expressed genes, absolute log2FC>=1 & q value <= 0.1) between infected versus uninfected in WT and *p62^KD^* macrophages at 0-, 24- and 48-.hpi **(H-I)** Pie charts showing functional classes of mitochondrial-associated DEGs at different time points in infected WT and *p62^KD^* macrophages.

Paradoxically, while the autophagy adaptors were expected to facilitate xenophagy, knocking them down reduced the bacterial CFU, implicating that these adaptors could be possibly helpful in *Mtb* growth in the host macrophages. Likewise, knocking down *p62/SQSTM1* or *OPTN* in another human monocytic cell line U937 also reduced intracellular bacterial load (Fig. S2B).

To gain further mechanistic insights into the role of autophagy adaptors in governing bacterial growth and survival inside macrophages, we focused on p62/SQSTM1 since it showed the highest recruitment to both mCherry-H37Rv phagosomes and TOM20^+^ mitochondria. We targeted p62/SQSTM1 by CRISPR/Cas9-mediated gene knockout in THP-1 macrophages (see methods) using gRNA designed against exon 3 of the primary protein-coding transcript (see methods). We achieved ∼80% knockdown using p62/SQSTM1 gRNA and hCAS9 co-transfection followed by selection as assessed by immunoblotting (Fig. S2C) and immunofluorescence (Fig. S2D). We continued using these stable cells as p62 knockdown (*p62^KD^*) cells and repeatedly analysed p62 expression to ensure efficient downregulation even after infection with *Mtb* (Fig. S2E). Knockdown of *p62/SQSTM1* did not overtly alter the cellular autophagy response since no discernible alterations in LC3-II levels were observed in the absence or presence of bafilomycin A_1_ (BafA_1_) (Fig. S2F).

Likewise, *p62/SQSTM1* knockdown did not affect the xenophagy response against H37Rv, as measured by colocalisation studies with LC3B (Fig. S2G). Similar to siRNA knockdown experiments, in *p62^KD^*THP-1 cells, bacterial CFU was consistently lower than in the WT cells (Fig. 2B). To confirm whether the reduction in bacterial CFU was due to slowing down of replication or enhanced killing in *p62^KD^* cells, we used the replication clock plasmid (21). The replication clock plasmid (pBP10) is unstable and contains the Kanamycin resistance gene that gets lost from *Mtb* during active replication. Loss of Kan^R^ during assay is indicative of active replication. A time-course experiment with pBP10 containing H37Rv clearly showed that the gradual loss of Kan^r^ over time in both WT and *p62^KD^* cells was indistinguishable, suggesting that the lower *Mtb* CFU observed in *p62^KD^* cells compared to the WT cells is due to a loss of survival and not due to a slowing down of replication (Fig. 2C).

Furthermore, we knocked down *p62* using siRNAs in human Monocyte-derived Macrophages (hMDMs) from three independent donors. In each case, we observed significant but varying degrees of decline in bacterial CFU in the knockdown cells (Fig. 2D). Thus, the absence of autophagy adaptor p62 appears to hamper *Mtb* growth and survival in THP-1 cells, U937 cells and also in the human primary macrophages, indicating it to be a general phenomenon in human macrophages.

In other animal models like zebrafish and mice, there are reports on p62/SQSTM1 and OPTN being host resistance factors against *Mtb* infection (22). We knocked down *p62/SQSTM1* and *OPTN* in the mouse BMDMs and did not observe any significant difference in *Mtb* CFU (Fig. S2H-I). Similarly, in a HOXB8-derived *p62^KO^* mouse monocytic cell line, there was no effect on *Mtb* CFU between WT and *p62^KO^* cells (Fig S2J). These results were consistent with another report, where bacterial CFU load in the lungs of p62 knockout mice was not significantly different from the wild-type animals (23). Therefore, autophagy adaptors like p62/SQSTM1 and OPTN may have some distinct species-specific functions in mouse and human macrophages.

### Perturbations in mitochondria-associated gene expression in *p62^KD^* cells upon *Mtb* infection

To understand the role of autophagy adaptors, specifically p62/SQSTM1, in regulating bacterial survival, we sought to elucidate the underlying mechanism behind the bacterial killing in the *p62^KD^* cells. In an unbiased approach, we performed gene expression analysis of WT and *p62^KD^* cells under uninfected and infected conditions at 0-, 24- and 48- hpi (methods). Overall, both WT and *p62^KD^*cells showed similar gene expression patterns across the three-time points and broadly similar expression patterns upon H37Rv infection.

However, the magnitude of regulation showed some differences between the WT and *p62^KD^* cells at later time points (Fig. S3A, Table S1). This was reflected in the gene ontology analysis. In both WT and *p62^KD^* cells, functional classes like a response to cytokine stimulus, inflammatory responses and response to LPS stimulation were equally upregulated (Fig. S3B). There were also some unique upregulated events in the *p62^KD^* cells, like NFκB pathway upregulation at 0 hpi, chemokine response pathways at 24 hpi and inflammatory responses at 48 hpi (Fig. S3B). In the cellular component GO class, most regulated genes belonged to trafficking vesicles, Golgi apparatus and Mitochondria (Table S2). Since p62/QSTM1 was the central autophagy adaptor getting recruited at the mitochondria, we specifically examined mitochondrial genes (Table S3). While at 0 hpi, WT and *p62^KD^* cells appeared almost identical in terms of gene expression, as the infection progressed, some differences emerged at later time points (24 and 48 hpi) (Fig. 2E-G). While genes like *RP2, GSR* and *OXR1* were upregulated only in the WT cells at 24 hpi, several other genes like *ETFBMKT, SLC25A19, SOD2, FTH1* and *BNIP3* were upregulated only in the *p62^KD^* cells (Fig. 2F). Increased *BNIP3* expression in the *p62^KD^* cells was exciting since it along with proteins like FUNDC1 and NIX constitutes ubiquitin independent autophagy adaptors and can directly tag damaged mitochondria for mitophagy-mediated degradation (24). Another interesting observation was the significantly enhanced expression of *MGME1,* a mitochondrial genome maintenance protein, in the *p62^KD^* cells at 48 hpi that gets downregulated upon *Mtb* infection (Fig. 2G). Overall, the mitochondria-targeted genes showing differential regulation upon *Mtb* infection at later time points included functional classes like lipid metabolism, mitochondrial nucleotide metabolism, immune response, redox regulation, mitophagy, autophagy and trafficking (Fig. 2H-I, Table S4). Thus, *p62^KD^*cells could exhibit a few differences in mitochondrial functioning or dynamics upon infection compared to the infected WT cells. In addition, *p62^KD^*cells showed dysregulated cytokine expression upon H37Rv infection compared to the WT cells (Fig. S3B, Table S5). Some of these cytokines, which consistently showed increased expression in *p62^KD^* cells compared to the WT cells, were IL1B, TNF, CXCL6 and CXCL2, suggesting a more inflammatory phenotype in the *p62^KD^* cells upon H37Rv infection. The results above indicate that p62/SQSTM1 is involved in mitochondrial dynamics and controlling inflammatory pathways during infection.

### Mitochondrial quality is maintained during H37Rv infection in p62/SQSTM1-depleted cells

The gene expression analysis suggested that the WT and *p62^KD^* cells responded differently to *Mtb* infection, particularly for genes involved in mitochondrial dynamics and function. However, how that could undermine mycobacterial growth and survival in those cells is unclear. Given the role of autophagy adaptors, specifically p62/SQSTM1 in the process of mitophagy (25–28) and also the fact that it was the maximally recruited autophagy adaptor on the mitochondria (Fig. 1F), we measured the parameters for assessing the mitochondrial function and dynamics in the cell. MitoTracker green (MTG) staining showed that there was a marginal increase in the total mitochondrial content in the *p62^KD^* cells upon H37Rv infection at early time points but not at the later time points (Fig. 3A). Moreover, there was no significant change in the overall status of mitochondrial depolarisation, as observed by TMRE staining at 0-, 24- and 48- hpi in *p62^KD^* and WT cells (Fig. 3B). Next, we measured mitochondrial superoxide that indicates the presence of active mitochondria, using MitoSox dye. While there was not much difference between the WT and *p62^KD^*cells, both cells, however, showed a tendency for an increase in the mitochondrial superoxide upon *Mtb* infection (Fig. 3C). The infection-dependent increase in MitoSox signal increased as the infection progressed from 0- to 24- and 48 hpi (Fig. 3C). However, only at the later time-point (48 hpi), this increase was significant in *p62^KD^* cells (Fig. 3C). The seahorse extracellular flux analyzer further examined mitochondrial respiration by performing the mitostress test, where we measured the cellular oxygen consumption rate (OCR) in response to the sequential addition of inhibitors of the electron transport chain (ETC). Overall, the OCR was found to be similar between WT and *p62^KD^* cells, although the latter showed a non-significant increase in basal and maximal respiration (Fig. 3D). Additionally, there was no significant difference in the mitochondrial DNA content and the expression of genes involved in mitochondrial fission/fusion in WT versus *p62^KD^* cells even after infection (Fig. S4A-B).

**Fig. 3.**
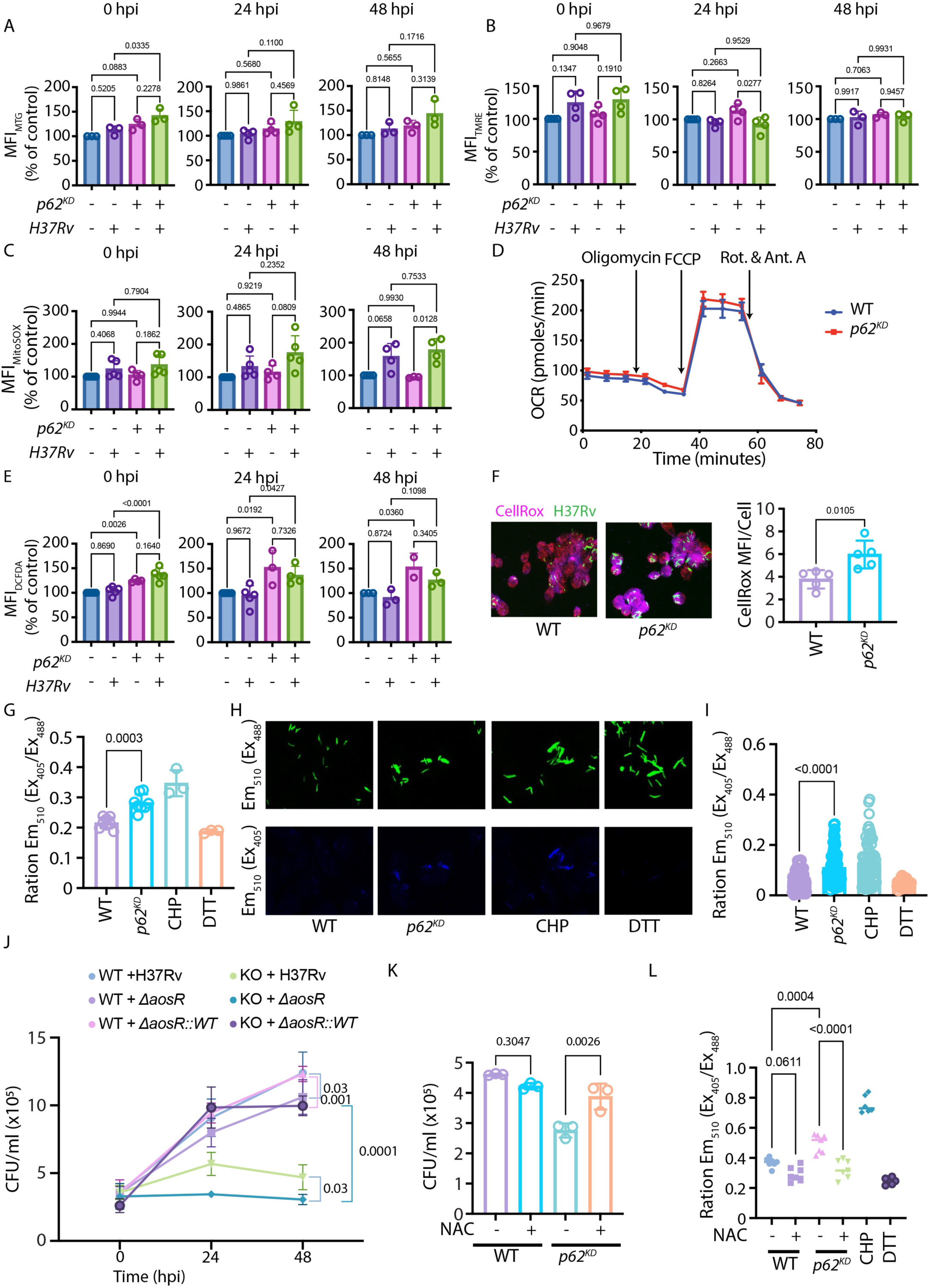
Mitochondrial quality is maintained during H37Rv infection in the absence of p62/SQSTM1: **(A-C)** The bar graphs represent the MFI (% of control) of (A) MTG (MitoTracker Green), (B) TMRE (Tetramethylrhodamine, ethyl ester), and (C) MitoSOX, via flow cytometry at 0-, 24- and 48-hpi in H37Rv-infected and uninfected WT and *p62^KD^* THP-1 macrophages. Data show mean ± SD, three independent experiments. **(D)** Mitostress (Seahorse extracellular flux assay) test in H37Rv-infected WT and *p62^KD^* macrophages at 24 hpi. Data show mean ± SD, three independent experiments. **(E)** DCFDA MFI (% of control) in WT and *p62^KD^* THP-1 macrophages at 0-, 24- and 48-hpi. Data show mean ± SD, three independent experiments. **(F)** IF analysis of CellROX staining (magenta) in WT and *p62^KD^* THP-1 macrophages at 24 hpi. The bar graph represents CellROX MFI. Data show mean ± SD, n=5 from two independent experiments. **(G)** The ratio of emission MFI of Mrx1-roGFP reporter at 510nm as excited at 405nm to 488nm via flow cytometry in WT and *p62^KD^* cells infected with the Mrx1-roGFP2 expressing H37Rv at 24 hpi. CHP and DTT were used as oxidising and reducing controls, respectively. Data show mean ± SD, three independent experiments. **(H-I)** Confocal microscopy images of H37Rv::Mrx1-roGFP2 in WT and *p62^KD^*cells, at 24 hpi, when excited at 405nm and 488nm. The bar graph depicts the ratio of emission MFIs at 510nm obtained upon excitation at 405nm to that at 488nm. Data show mean ± SD, n>20 fields, 30-50 bacteria per field, from two independent experiments. **(J)** Time course CFU analysis of H37Rv*ΔaosR* and complemented strain H37Rv*ΔaosR::WT* in WT and *p62^KD^*cells. Data show mean± SD, representative of two independent experiments **(K)** CFU analysis of H37Rv in the presence of N-acetyl cysteine (NAC, 2mM for 24 hours) in *p62^KD^* macrophages at 24 hpi. Data show mean ± SD, three independent experiments. **(L)** Dot plot depicts the ratio of emission MFI of *Mrx1-roGFP2*::*H37Rv* at 510nm as excited at 405nm to 488nm via confocal microscopy in NAC treated and untreated WT and p62*^KD^* at 24 hpi. Data show mean ± SD, n>=6 fields, 30-50 bacteria per field, from two independent experiments.

Our data suggest that p62/SQSTM1 depletion does not overtly alter the mitochondrial function and dynamics, like content and mitochondrial outer membrane permeabilization (MOMP). However, we did observe a delayed increase in the mitochondrial respiration in *Mtb-*infected *p62^KD^* cells, indicating involvement and/or activation of specific quality control pathways in these cells.

### *Mtb* witnesses increased redox stress in *p62^KD^* cells

While the usual mitochondrial quality parameters were unperturbed in the *p62^KD^* cells upon H37Rv infection, an infection-induced increase in the production of mitochondrial superoxide indicated the possibility of a redox imbalance in these cells. Indeed, there was a significant increase in the reactive oxygen species (ROS), as measured by DCFDA staining in the *p62^KD^*macrophages in the uninfected state or upon H37Rv infection compared to the corresponding WT conditions (Fig. 3E). Increased ROS in the infected *p62^KD^* cells was also corroborated with significantly higher CellRox fluorescence observed in the *Mtb* infected *p62^KD^*cells compared to the WT cells (Fig. 3F). Next, we examined, whether *Mtb* too experienced a more oxidising environment in *p62^KD^* cells and if that contributed to their diminished growth in these cells. To do this, we utilized a genetically encoded biosensor of mycothiol redox potential in *Mtb* (*mrx1-roGFP2*) (29). The dynamic changes in mycothiol redox potential serve as a proxy for changes in the cytoplasmic redox state of *Mtb*. The redox state of *Mtb* was quantified by measuring the Mrx1-roGFP2 fluorescence ratio (excitation: 405 nm/488 nm; emission: 510 nm). Upon oxidation, the biosensor ratio increases, whereas the ratio decreases in response to the reductive stress (30). Flow cytometry and imaging data revealed that *Mtb* was exposed to a highly oxidising environment in the *p62^KD^* cells compared to the WT cells (Fig. 3G-I). To test whether increased oxidative stress impacted bacterial growth in these cells, we used a redox-sensitive strain of *Mtb*- *H37RvΔaosR*. The *H37RvΔaosR* lacks a critical cysteine biosynthesis pathway, thereby rendering the strain sensitive to oxidative stress (31). Similar to the WT H37Rv strain, the H37Rv*ΔaosR* strain showed a significant decline in CFU in the *p62^KD^* cells compared to the WT THP-1 cells (Fig. 3J). Interestingly, the CFU count of redox-sensitive strain H37Rv*ΔaosR* was significantly lower than the WT H37Rv strain in *p62^KD^* cells (Fig. 3J). Moreover, complementation of the deleted strains with WT *aosR* rescued the bacterial CFU in the *p62^KO^*cells (Fig. 3J). While the *ΔaosR::WT* strain showed higher CFU in the *p62^KO^* cells than the H37Rv, it was significantly lower than the *aosR::WT* CFU in the WT cells (Fig. (3J). These results suggest that the elevated oxidative stress witnessed by *Mtb* may contribute to a decline in its CFU count in *p62^KD^* cells. To test this hypothesis, we used the ROS scavenger N-acetyl cysteine (NAC), which significantly rescued the bacterial CFU in the NAC-treated *p62^KD^* cells compared to the untreated *p62^KD^* cells (Fig. 3K). Using *mrx1-roGFP2* reporter, via confocal microscopy we verified that NAC treatment indeed led to a decrease in the level of oxidative stress witnessed by *Mtb* in the *p62^KD^* cells. (Fig. 3L) These data suggest that the depletion of p62/SQSTM1 in THP-1 macrophages enhances redox stress, contributing to reduced mycobacterial growth and survival in these cells.

### Understanding mitochondrial quality control in *p62^KD^* cells

We next studied if mitochondria contributed to the enhanced cellular ROS in *p62^KD^* cells, despite showing normal mitochondrial parameters. Mitochondria may act as the direct source of ROS for bacterial killing upon their recruitment to the bacterial phagosome (32). In *p62^KD^* cells, there was significantly enhanced recruitment of the mitochondria to the bacterial phagosomes, as evident from mitochondrial marker protein TOM20 (translocase of outer mitochondrial membrane 20) immunostaining and 3D-reconstructions to analyse TOM20 recruitment on H37Rv structures (Fig. 4A). Since TOM20 recruitment to *Mtb* correlated with reduced bacterial growth, we suspected that the phagosome-recruited mitochondria could contribute to the bacterial killing. We also hypothesised that the fraction of mitochondria recruited to the phagosomes could be mostly depolarised, considering they are the major source of mitochondria-generated ROS (33, 34). The ubiquitin ligase Parkin is known to get stabilised on the depolarised mitochondria and ubiquitinates mitochondrial outer membrane proteins for mitophagy targeting (35, 36). Contrary to our expectation, there was a decline in Parkin-TOM20 co-localization in *p62^KD^* cells (Fig. 4B). However, unlike Parkin, there was an increase in the ubiquitin-TOM20 interaction in the *p62^KD^* cells (Fig. 4C). Curiously, the pattern of Ubiquitin and Parkin recruitment to TOM20 mirrored with their recruitment to H37Rv, where there was an increase in the Ubiquitination of *Mtb* but a decline in Parkin recruitment (Fig. 4D-E). The contrasting observation of enhanced ubiquitination and reduced Parkin recruitment at TOM20^+^-mitochondria and *Mtb*-containing phagosomes was intriguing. Since ubiquitinated mitochondria could undergo autophagy-mediated degradation or mitophagy, we next checked the status of mitochondrial localization of LC3B in *p62^KD^*cells upon infection. We analysed the co-localization of TOM20 with LC3B in the WT and *p62^KD^* cells. There was a significant increase in TOM20-LC3B colocalisation in *p62^KD^* cells compared to the WT cells (Fig. 4F). We also included a BafA_1_ treatment group for both WT and *p62^KD^*cells in this assay to examine whether LC3B^+^-mitochondria positive vesicles matured and fused with the lysosomes distinctly in the *p62^KD^* cells compared to the WT cells. We expected an increase in TOM20-LC3B colocalization in the BafA_1_-treated group compared to the corresponding untreated controls, as BafA_1_ would block both LC3B and TOM20 degradation. Surprisingly, in either the WT or *p62^KD^* cells, BafA_1_ treatment did not increase the TOM20-LC3B colocalisation (Fig. 4F). However, BafA_1_-treated *p62^KD^* cells had significantly higher TOM20-LC3B colocalisation compared to the corresponding WT cells. Thus, whereas more mitochondrial cargos were targeted towards LC3B^+^-compartments in the *p62^KD^* cells, the mitochondrial degradation was less efficient in both WT and *p62^KD^* cells.

**Fig. 4.**
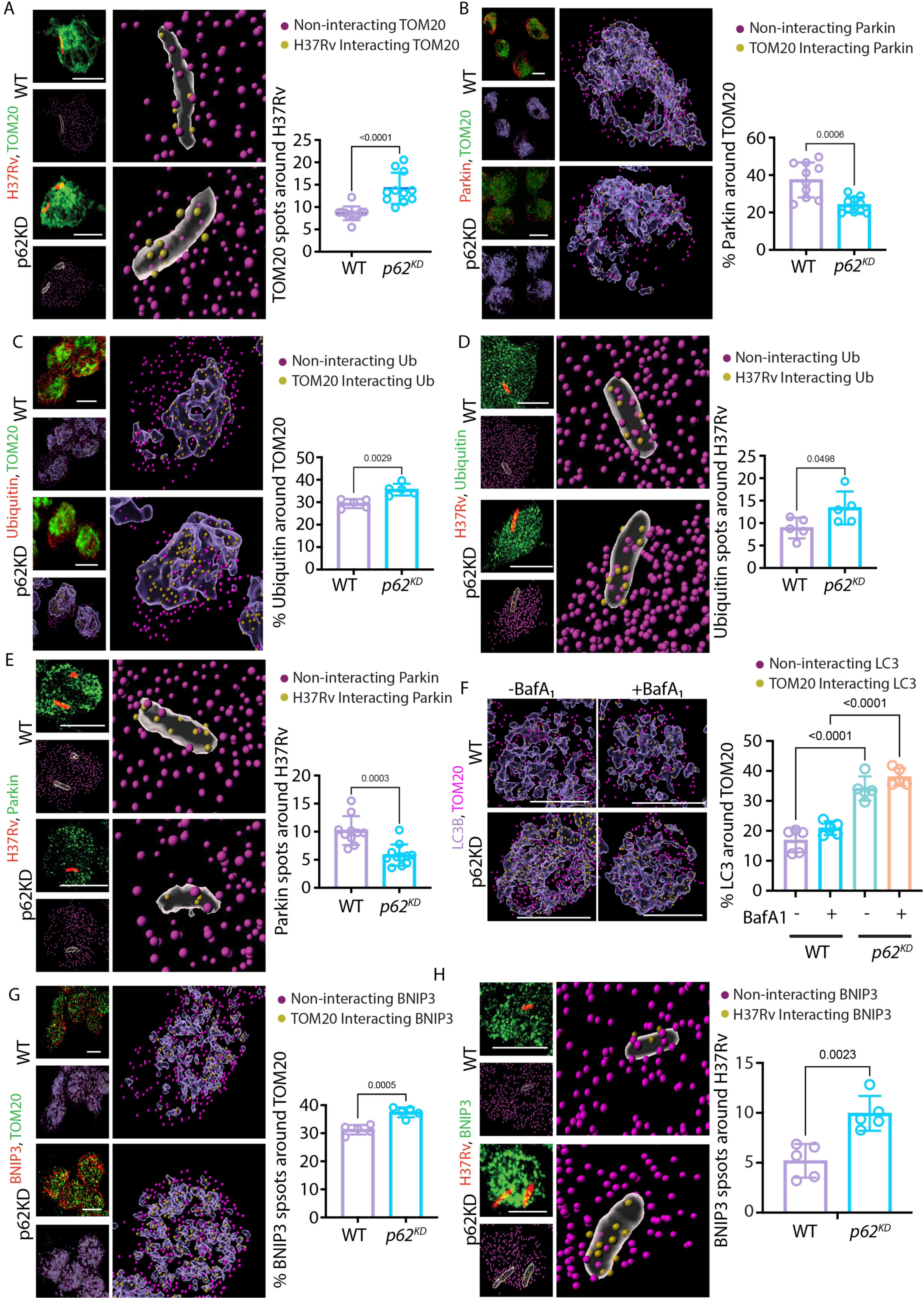
Mitochondrial Quality Control in p62/SQSTM1 KD cells: **(A)** Immunofluorescence images for mCherry-H37Rv (Red) and TOM20 (green) interaction in WT and *p62^KD^*THP-1 macrophages at 24 hpi. The 3D reconstruction of H37Rv structures and TOM20 spots is shown at the right; the bar graph shows the average number of yellow TOM20 spots/bacteria. Data show mean ± SD, n> 10 fields, 20-35 cells per field, from three independent experiments, **(B-C)** Immunofluorescence images of TOM20 (green) and Parkin (red, B) or Ubiquitin (red, C) respectively. Representative TOM20 3D structures and Parkin or Ubiquitin spots are shown on the right. The bar graph shows per cent of Ubiquitin (B) or Parkin (C) spots interacting with the TOM20 structures. Data show mean± SD, n>6 fields, 20-35 cells per field, from two independent experiments. **(D-E)** Immunofluorescence images of Ubiquitin (D) and Parkin (E) respectively with mCherry-H37Rv, and 3D reconstructed images of H37Rv structures and Ubiquitin (D) or Parkin (E) spots. The bar graph shows per cent of spots <0.2 μm from *Mtb*. Data show mean ± SD, n≥5 fields, >20-35 cells per field, from two independent experiments. **(F)** Mitophagy flux assessed by the interaction of LC3B spots and TOM20 structures in the presence and absence of bafilomycin A1 (BafA_1_, 100nM, 3 hours) in WT and *p62^KD^* THP1 macrophages at 24 hpi. Data show mean ± SD, n≥6 fields, 20-35 cells per field, from two independent experiments. **(G-H)** Immunofluorescence images on the left show the interaction of BNIP3 spots with TOM20 (G) or H37Rv (H) structures, respectively. The bar graphs depict the number of BNIP3 spots <0.2 μm from TOM20 (G) or H37Rv (H) structures. Data show mean ± SD, n=5 fields, 20-35 cells per field, from two independent experiments. For all 3D reconstructions, yellow spots <0.2 μm from TOM20 or *Mtb* structures, magenta spots >0.2 μm from TOM20 or *Mtb* structures; Scale bar: 10 μm.

Since similar BafA1 treatment shows increased LC3II upon immunoblotting (Fig. S2F), we infer that autophagic degradation of cargos other than mitochondria continues in *p62^KD^* cells. Could an increase in ubiquitination alone explain the enhanced LC3B recruitment to TOM20 in p62-depleted cells? LC3B can also be recruited to mitochondria through ubiquitin-independent autophagy receptors like BNIP3, FUNDC1, NIX, etc. Incidentally, BNIP3, but not FUNDC1 or NIX, was also among the highly upregulated genes in *Mtb*-infected *p62^KD^* cells (Fig. 2F), a finding that we confirmed at the protein level by immunoblotting (Fig. S4C). The increased BNIP3 levels in the *p62^KD^* cells were also associated with a significantly increased BNIP3 recruitment to both TOM20 and *Mtb* as compared to the WT cells (Fig. 4G-H).

Greater recruitment of BNIP3 and also TOM20, both of which are mitochondrial outer membrane proteins, to *Mtb*-phagosomes indicated that in *p62^KD^* cells, damaged mitochondria or possibly mitochondria-derived vesicles (MDVs) could be recruited to *Mtb*-phagosomes on their way to being targeted for mitophagy. While significantly higher BNIP3 recruitment, as well as ubiquitination, could explain enhanced LC3B recruitment to TOM20, overall, the pattern of recruitment of several key molecules to mitochondria and *Mtb* phagosomes in *p62^KD^*cells was incongruent with the anticipated mitophagy pathways and indicated that there were additional mechanisms operational to ensure MQC in *p62^KD^* cells. Results so far, while strongly suggesting crosstalk between mitochondria and *Mtb* phagosomes, were inconclusive regarding the nature of mitochondrial recruitment to the *Mtb* phagosomes and whether and how that contributed to the enhanced redox stress to *Mtb* in the *p62^KD^* cells.

#### H37Rv-infected *p62^KD^* cells show enhanced biogenesis of TOM20^+^-Mitochondria-derived vesicles (MDVs)

While mitophagy is the primary mechanism for MQC in diverse cell types, increasingly, MDVs are being reported as an important mechanism for MQC, specifically in those cells where mitophagy is compromised (37, 38). MDVs are small vesicles (50-200 nm in size) derived exclusively from the outer or inner mitochondrial membrane that buds off from the damaged parts of mitochondria and fuses with the lysosome for autophagy-independent degradation (39, 40). Analysis of TOM20 immunostaining in *p62^KD^* THP-1 cells under high-resolution microscopy revealed a large number of smaller TOM20^+^ vesicles-like structures in a size range of 200 nm or smaller compared to the WT cells (Fig. 5A). Dual staining with TOM20 and MCU, which respectively are mitochondrial outer membrane (OM) and inner membrane (IM) proteins, strongly indicated that indeed the smaller TOM20 structures seen in *p62^KD^* cells represented MDVs. While we could notice both IM-derived (TOM20^-^MCU^+^) and OM-derived (TOM20^+^MCU^-^) MDVs (Fig. 5B) in WT and *p62^KD^* cells, we counted the number of TOM20^+^ vesicles in the size range of 100-200nm and observed more than two-fold increase in the number of these vesicles in *p62^KD^* cells compared to the WT cells (Fig. 5C). Interestingly, *Mtb* infection did not alter the MDV numbers in the WT or *p62^KD^* cells further (Fig. 5C), suggesting that MDV biogenesis reflected an infection-independent mechanism, which most likely was activated as an MQC pathway in the absence of autophagy adaptor p62/SQSTM1. To further confirm that in *Mtb*-infected *p62^KD^* cells, there was an increase in the formation of TOM20^+^-MDVs, we analysed TOM20 co-localization with additional markers from the mitochondrial IM (MCU and ECSIT) or the mitochondrial matrix (PDHA1, HSP60, SOD2) (Fig. 5D). We anticipated that TOM20^+^ MDVs being exclusively derived from the OM of the mitochondria should show selectivity in terms of its association with IM proteins and matrix proteins. Among the three mitochondrial matrix proteins studied, only SOD2 showed a marginal increase (from ∼66% in the WT cells to ∼73% in *p62^KD^* cells) in its association with TOM20 in the *p62^KD^* cells compared to the WT cells (Fig. 5E). In contrast, PDHA1 and HSP60 showed no significant difference (Fig. 5E, S5A-C). Unlike the matrix proteins, there was a substantial decline in TOM20 colocalisation with the mitochondrial IM proteins ECSIT and MCU (Fig. 5F and S5D-E). The clear distinction in the association between TOM20 and IM proteins (ECSIT and MCU) and OM proteins (BNIP3, Fig. 4), coupled with the increased TOM20^+^-vesicles observed in *p62^KD^* cells, further suggested an increase in OM-derived MDV-biogenesis in *p62^KD^*cells. Taken together from the results in the above sections, it was evident that p62-depleted macrophages trigger TOM20^+^-MDV biogenesis.

**Fig. 5.**
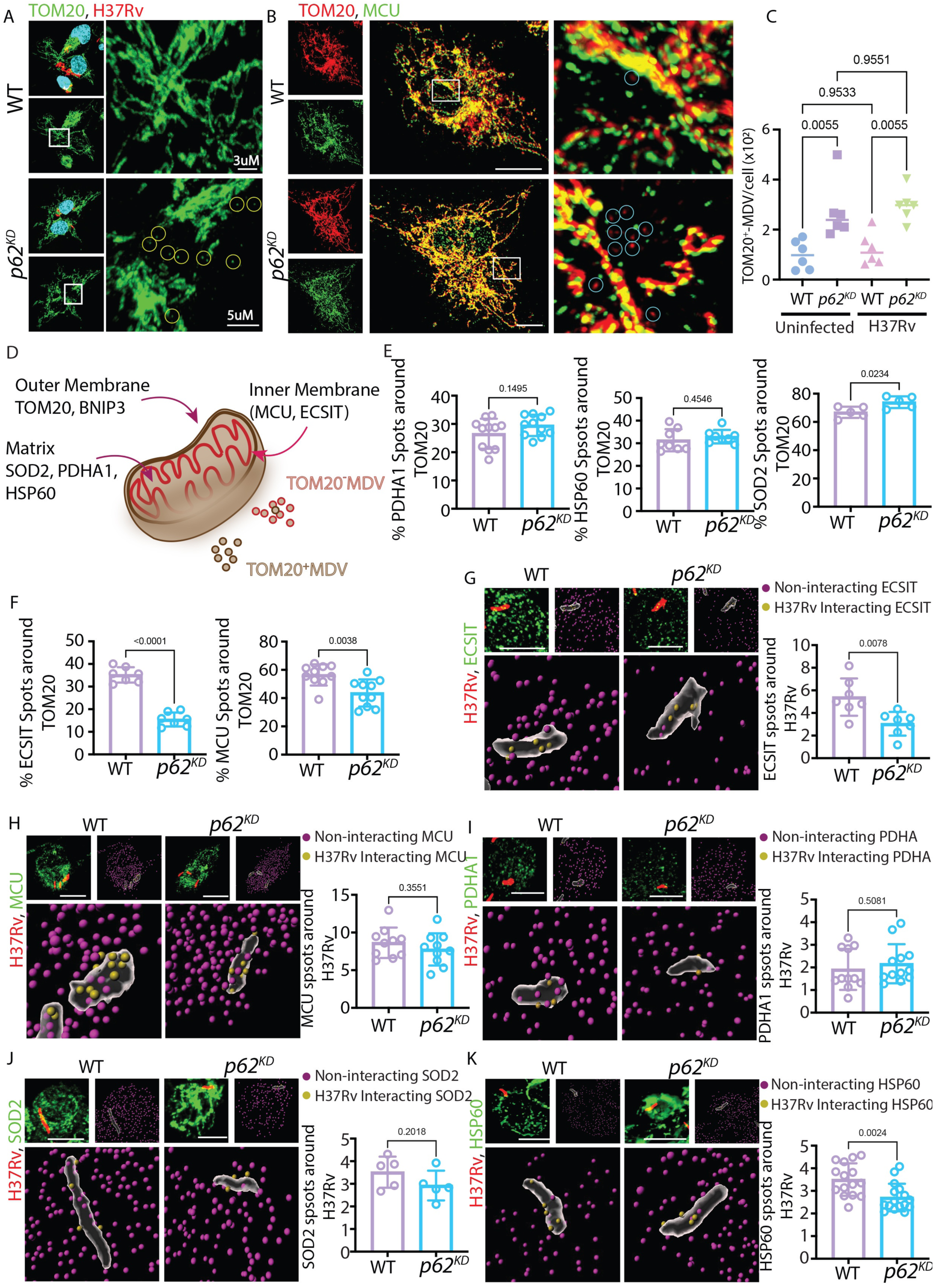
Enhanced TOM20^+^-Mitochondria-derived vesicles (MDVs) biogenesis in *p62^KD^* cells: **(A)** High-resolution images of immunostained TOM20 (green) in H37Rv-infected WT and *p62^KD^*macrophages at 24 hpi. The zoomed images on the right show encircled TOM20^+^-MDVs in the *p62^KD^* cells. **(B)** High-resolution images of immunostained TOM20 (red) and MCU (green). The encircled area depicts TOM20^+^MDVs in the zoomed images at the right. **(C)** Quantification of number of TOM20^+^MDVs in uninfected and *Mtb*-infected WT and *p62^KD^* cells. Data show mean ± SD, n=6 fields having 5-10 cells per field, from two independent experiments **(D)** Schematic depiction of the presence of different proteins on mitochondrial matrix, inner and outer membrane. **(E)** Bar graphs show quantitative interactions of matrix mitochondrial protein spots of PDHA1, HSP60 and SOD2 with TOM20 structures. **(F)** Bar graphs show the quantitative association of inner membrane mitochondrial protein spots of ECSIT and MCU with TOM20 structures. (**G**-**K**) Immunofluorescent images depict the interaction of mCherryH37Rv (red) with ECSIT **(G),** MCU **(H**), PDHA1 **(I)**, SOD2 **(J)** and HSP60 **(K)** (green), respectively, in WT and *p62^KD^* THP-1 macrophages at 24 hpi. The 3D reconstruction shows H37Rv structures and indicates mitochondrial protein spots. The yellow spots are <0.2 μm from H37Rv structures, and the magenta spots are >0.2 μm from bacterial structures. The bar graph on the right shows a number of mitochondrial protein spots on *Mtb*. Data show (E-K) mean ± SD, n≥5 fields having 20-35 cells per field, from two independent experiments, Scale bar =10 μm unless specified.

### TOM20 recruited at *Mtb* phagosomes in *p62^KD^* cells represent TOM20+-MDVs

Given the results above, it was imperative to assess whether the enhanced TOM20 recruitment on *Mtb* phagosomes in *p62^KD^* cells was actually due to the recruitment of TOM20^+^-MDVs in these cells. We examined the recruitment of mitochondrial IM and matrix markers shown in Fig. 5E-F for their recruitment to *Mtb* phagosomes. There was a significant decline in the recruitment of mitochondrial IM marker ECSIT to *Mtb*-phagosomes in the *p62^KD^* cells, a pattern similar to ECSIT-TOM20 colocalisation (Fig. 5G, S5D). Unlike ECSIT, there was no decline in the recruitment of the other IM protein MCU on *Mtb*-phagosomes (Fig. 5H, S5E). Since MCU^+^TOM20^-^ MDVs are also formed in both WT and *p62^KD^* cells (Fig. 5B), it is possible that some of the MCU^+^MDVs continue getting recruited to the phagosomes in both WT and *p62^KD^* cells. Additionally, it is possible that ECSIT proteins were not included in the MCU^+^ IM-derived MDVs. The matrix proteins PDHA1, SOD2, and HSP60 did not entirely follow the pattern for *Mtb* colocalisation vis-à-vis TOM20 colocalisation (Fig. 5I-K, SA-C). Overall, these colocalisation studies reveal that TOM20 and *Mtb* showed a remarkable similarity in increased recruitment of OM (BNIP3) and decreased recruitment of IM (ECSIT) proteins. However, the matrix proteins did not show a strong correlation. These results suggest that in *p62^KD^* cells, there is a significant increase in OM-derived TOM20^+^-MDVs formation and that a large fraction of these MDVs gets recruited to the *Mtb*^+^-phagosomes upon infection.

#### TOM20^+^-MDVs overcome phagosome maturation arrest imposed by *Mycobacterium tuberculosis*

We next investigated whether MDV recruitment to the *Mtb*-phagosomes contributed to bacterial growth and survival phenotype observed in these cells. MDVs, once formed, can either be secreted out of the cells as extracellular vesicles (EVs) or fuse with intracellular compartments, such as lysosomes, where they are degraded (41–43). Since TOM20 levels on *Mtb*-phagosomes were increased in the *p62^KD^* cells, it is possible that TOM20^+^-MDVs were recruited to the *Mtb*-phagosomes while en route to being targeted to the lysosomes. The lysosomal targeting of MDVs requires proteins from the Sortin-Nexin family, like SNX9 and the small GTPase RAB7 (37, 44). Incidentally, the exclusion of RAB7 from mycobacterial phagosomes and autophagosomes is a widely studied mechanism that helps the bacteria evade the lysosomal targeting (45). In *p62^KD^* THP-1 cells, there was a significant increase in RAB7 recruitment to *Mtb*-phagosomes compared to the WT cells (Fig. 6A). Increased RAB7 recruitment on the bacterial phagosomes should also result in increased lysosomal targeting of *Mtb* in these cells. In agreement with that, there was a significant increase in mycobacterial localisation to LysoTracker, Cathepsin D, and LAMP1 positive compartments in *p62^KD^*cells (Fig. 6B-D). It is, therefore, possible that TOM20^+^-MDVs, while getting recruited to the *Mtb* phagosomes for lysosomal targeting in *p62^KD^* cells, also help recruit RAB7, which unblocks the maturation arrest imposed by *Mtb* on these phagosomes. This was further corroborated by an increase in LAMP1-TOM20 colocalisation in *p62^KD^*cells, confirming the targeting of MDVs to the lysosomes (Fig. 6E). To verify if indeed RAB7 recruitment was involved in enhanced lysosomal targeting and impacted *Mtb* survival in *p62^KD^* cells, we knocked down *RAB7* using siRNA (Fig. S5F). In the WT cell, the knockdown of *RAB7* did not have much impact on H37Rv survival (Fig. 6F), which is expected since very little *RAB7* is known to get recruited to *Mtb* under WT conditions (45). However, the knockdown of *RAB7* in *p62^KD^* cells substantially improved *Mtb* growth and survival, suggesting that RAB7-recruitment indeed played a role in combating *Mtb* growth in these cells (Fig 6F). In agreement with this notion, the knockdown of RAB7 in *p62^KD^*cells decreased lysosomal targeting of *Mtb,* as evident from lower colocalisation of *Mtb* with LAMP1 (Fig. 6G). Curiously, RAB7 knockdown in the *p62^KD^* cells also led to a decline in the TOM20 recruitment on *Mtb* phagosomes, suggesting that TOM20^+^-MDV recruitment to *Mtb* phagosomes was somewhat dependent on RAB7 (Fig. 6H).

**Fig. 6.**
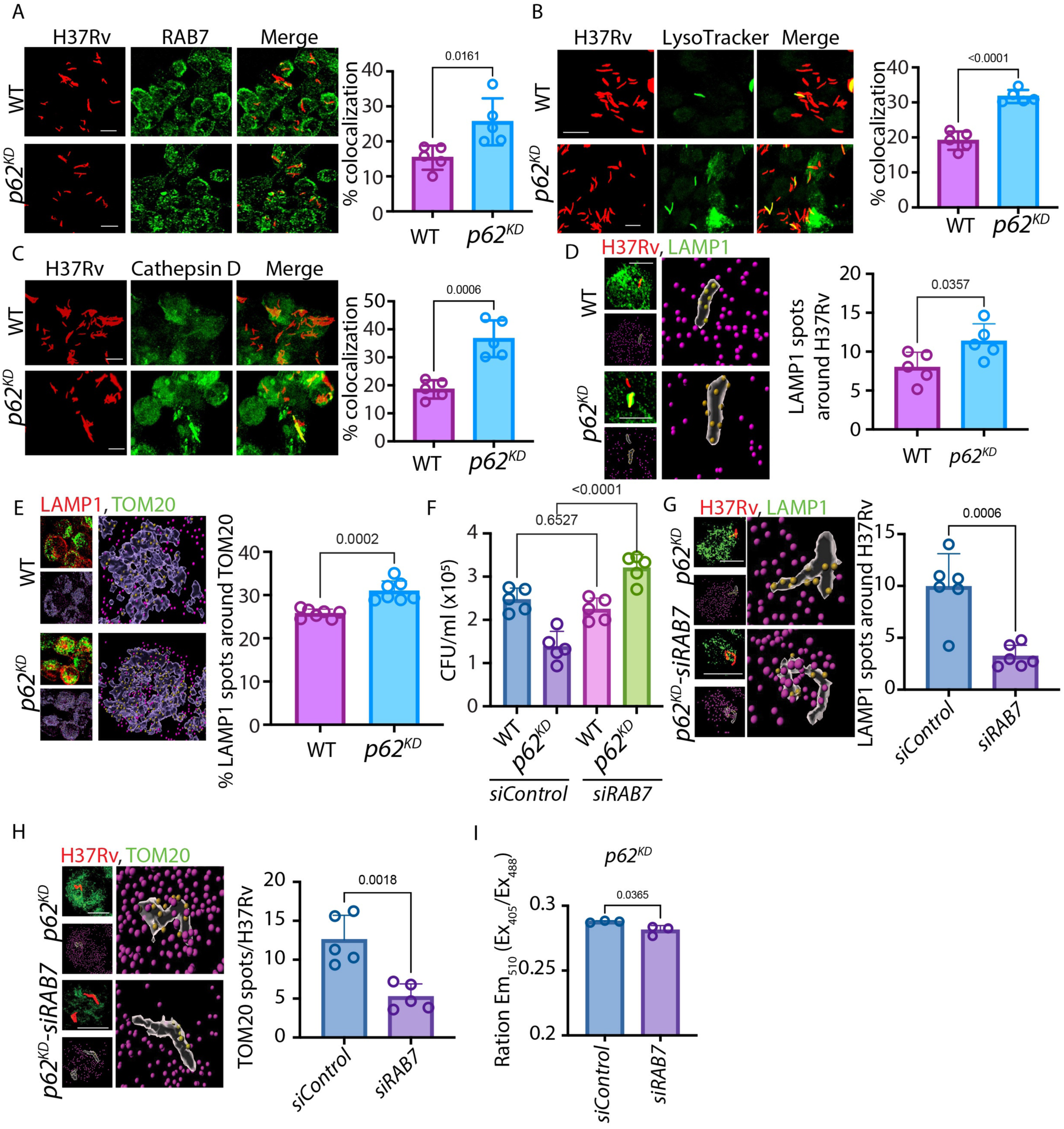
TOM20^+^-MDVs overcome phagosome maturation arrest imposed by *Mycobacterium tuberculosis*: **(A-C)** Confocal images show the colocalisation of mCherry H37Rv (red) with RAB7, Lysotracker and Cathepsin D respectively (green) in WT and *p62^KD^*THP-1 macrophages at 24 hpi. The bar graph shows per cent colocalisation, calculated manually using Imaris software. Data show mean ± SD, n=5 fields, 20-35 cells per field, from two independent experiments. **(D)** The image shows the interaction of LAMP1 spots with TOM20 structures. The bar graph depicts the number of LAMP1 spots <0.2 um (yellow) from the bacterial structure. Data show means ± SD, n=5 fields, 20-35 cells per field, from two independent experiments. **(E)** Images show the interaction of LAMP1 spots with TOM20 structures in *Mtb*-infected WT and *p62^KD^* cells. The plot shows LAMP1 spot quantification on TOM20 structure in *Mtb*-infected WT and *p62^KD^* cells. Data show mean ± SD, n> 5 fields, 20-35 cells per field, from two independent experiments **(F)** H37Rv CFU in WT and *p62^KD^* cells upon RAB7 knockdown at 24 hpi. Data show means ± SD, n=5. **(G-H)** Quantifying LAMP1 (G) and TOM20 (H) spots on H37Rv in RAB7 knockdown *p62^KD^* cells at 24 hpi. Data show means ± SD, n=5 fields, > 20 cells per field, from two independent experiments. **(I)** The relative oxidation state of *Mrx1-roGFP2* reporter as the ratios of emission MFI at 510nm when excited at 488nm to that of 405nm in *p62^KD^* and *RAB7* depleted *p62^KD^* cells. Data show means ± SD, n=5 Fields, > 40 bacteria per field, from two independent experiments. Scale bar: 10 μm

While it was now evident that *Mtb* was driven towards the lysosomes by TOM20+MDVs, we also witnessed ROS-mediated killing of *Mtb* in the *p62^KD^* cells (Fig. 3). We hypothesised that the increased oxidative stress seen by *Mtb* in *p62^KD^* cells could also be attributed to the recruitment of TOM20^+^-MDVs. At least one recent study shows similar MDV-mediated oxidative stress and killing of *Staphylococcus sp.* Bacteria (46). Since RAB7 knockdown in *p62^KD^* cells decreased TOM20^+^MDV recruitment on *Mtb* phagosomes, using H37Rv::*mrx1-roGFP2* bacteria, we found that the loss of TOM20^+^-MDV recruitment reversed the oxidative stress witnessed by *Mtb* in the *p62^KD^* cells (Fig. 6I). Our data implicate that TOM20^+^MDV recruitment on *Mtb* phagosomes could have anti-bacterial functions due to a combination of both redox stress and reversal of PMA.

#### MQC pathways intricately impact anti-bacterial responses in the macrophages

As evident from our data, TOM20^+^-MDV-dependent facilitation of lysosomal targeting of *Mtb* in *p62^KD^* cells showed a unique perspective in the crosstalk between mitochondrial homeostasis and cellular anti-bacterial responses. It is also evident that increased TOM20^+^MDVs observed in *p62^KD^* cells could be helping MQC in these cells since we did not observe any significant loss of MQC in the *p62^KD^*cells upon *Mtb* infection (Fig. 3). We therefore next studied whether the loss of MQC under these circumstances could further aid to the bacterial killing. Parkin is an important mediator of MQC, specifically since it triggers ubiquitin-mediated mitophagy and is also implicated in MDV biogenesis under diverse conditions (47, 48).

Expectedly, the knockdown of *PRKN* in *p62^KD^* THP-1 cells led to a loss of MQC, as was evidenced by a significant increase in mitochondrial depolarisation in *p62^KD^* cells (Fig. 7A). We also observed round, spherical TOM20 structures, typical of depolarised mitochondria upon *PRKN* knockdown in *p62^KD^* cells (Fig. 7B). Since the presence of depolarised mitochondria could cause more oxidative damage to the cell, we investigated the oxidative stress witnessed by *Mtb* in *p62^KD^*cells upon *PRKN* knockdown. We observed a slight but significant increase in the redox stress witnessed by *Mtb* in *p62^KD^* cells upon PRKN knockdown compared to *p62^KD^* cells alone (Fig. 7C). To test whether the loss of MQC in *p62^KD^* cells upon *PRKN* knockdown had any impact on intracellular bacterial growth and survival, we investigated *Mtb* CFUs in these cells. We observed significantly reduced *Mtb* survival in *PRKN* knockdown *p62^KD^* cells compared to the *p62^KD^*cells (Fig. 7D). *PRKN* knockdown did not impact MDV biogenesis as inferred from the absence of any difference in the recruitment of TOM20^+^-MDVs to the *Mtb* phagosomes (Fig. 7E-F). In agreement to that, there was no change in the lysosomal targeting of *Mtb* upon *PRKN* knockdown in *p62^KD^* cells (Fig. 7G). Thus, enhanced redox stress due to mitochondrial depolarisation, along with continued lysosomal targeting, caused excessive *Mtb*-killing in these cells. This was further corroborated by using Nigericin, a K^+^/H^+^ ion exchanger, which causes mitochondrial depolarisation (49). Treatment with nigericin led to a dramatic loss of mitochondrial outer membrane permeability (Fig. S5H) and showed significantly increased oxidation of Mrx1-RoGFP2 reporter (Fig. S5I). Moreover, similar to the results in *PRKN* knockdown in *p62^KD^* cells, nigericin treatment also led to a significantly reduced *Mtb* survival in the *p62^KD^* cells (Fig. S5J). At the doses used, nigericin did not impact host cell survival (Fig. S5K).

**Fig. 7:**
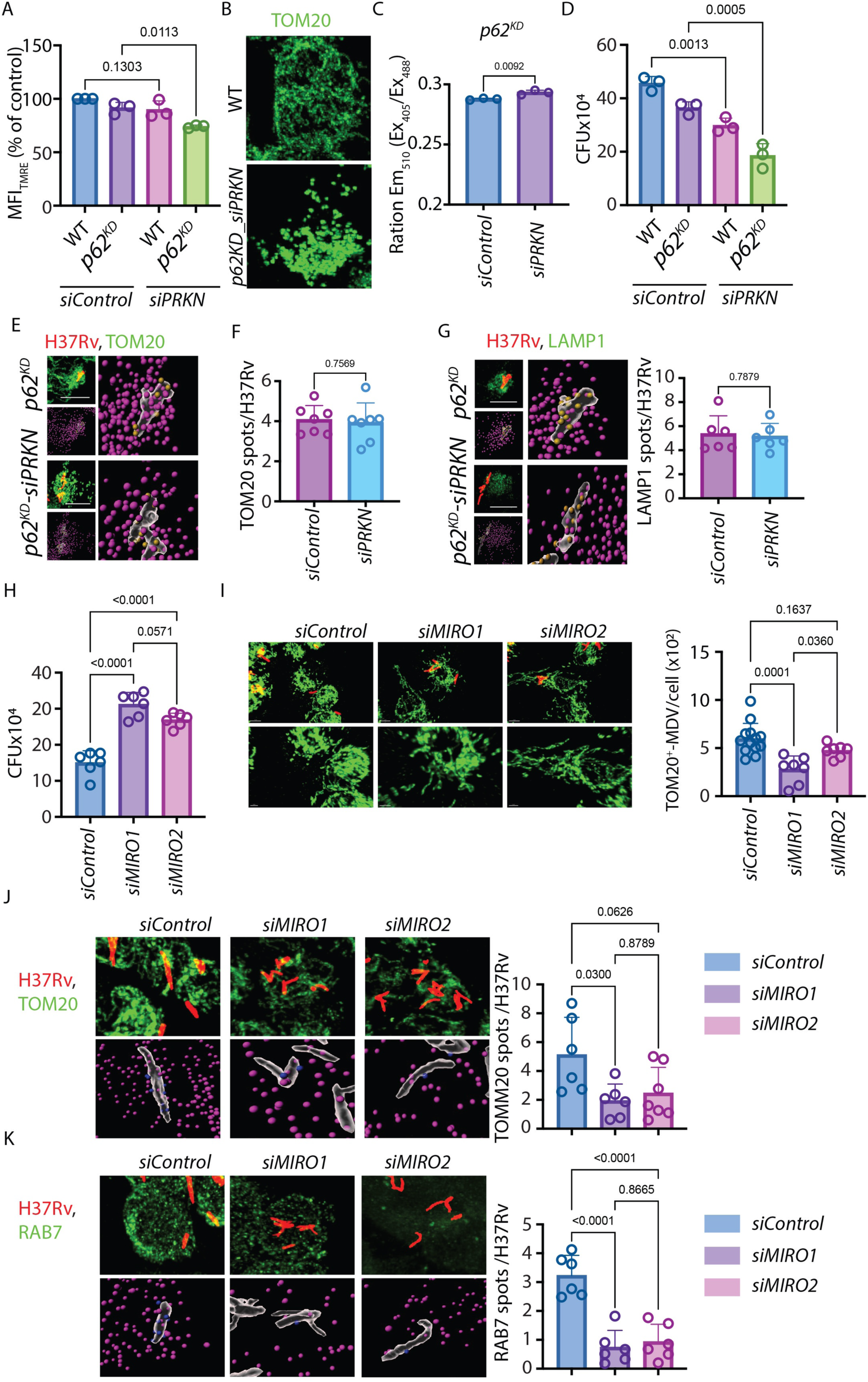
Triggering loss of MQC in *p62^KD^* cells aggravates bacterial killing. **(A)** TMRE per cent MFI in *PRKN* knockdown cells in WT and *p62^KD^*THP-1 macrophages. Data show mean ± SD from three independent experiments **(B)** The confocal images depict the circular depolarised mitochondria (immunostained with TOM20, green) in *PRKN*-depleted *p62^KD^*cells. **(C)** MFI ratio of *Mrx1-roGFP2*::*H37Rv* at 405nm to 488nm in *p62^KD^* and *Parkin-depleted p62^KD^* cells. Data show mean ± SD, n=5 Fields, > 40 bacteria per field, from two independent experiments. **(D)** H37Rv CFU in WT and *p62^KD^* THP-1 macrophages upon *PRKN* depletion at 24 hpi. Data show mean ± SD from three independent experiments. **(E-F)** TOM20 spots on H37Rv structures in *p62^KD^* cells upon *PRKN* knockdown. Data show means ± SD, n=7 fields, each having 20-35 cells, from two independent experiments. **(G)** LAMP1 spots on H37Rv structures in *p62^KD^* cells upon *PRKN* knockdown. Data show means ± SD, n=6 fields, 20-35 cells per field, from two independent experiments. **(H)** H37Rv CFU in *siCtrl*, *siMIRO1* and *siMIRO2* treated *p62^KD^* THP-1 macrophages at 24 hpi. Data show mean ± SD from two independent experiments. **(I)** High-resolution images of immunostained TOM20 (green) in *siCtrl*, *siMIRO1* and *siMIRO2* treated *p62^KD^* THP-1 macrophages at 24 hpi. The plot depicts the quantification of TOM20^+^ MDV in these cells. Data show means ± SD, n≥7 fields, 5-10 cells per field, from two independent experiments. **(J)** The images show the interaction of TOM20 spots with H37Rv structures in *siCtrl*, *siMIRO1* and *siMIRO2* treated *p62^KD^* THP-1 macrophages at 24 hpi. The bar graph depicts the number of TOM20 spots <0.2 μm (blue) from the bacterial structure. Data show means ± SD, n=6 fields, 20-35 cells per field, from two independent experiments. (K) The image shows the interaction of RAB7 spots with H37Rv structures in *siCtrl*, *siMIRO1* and *siMIRO2* treated *p62^KD^*THP-1 macrophages at 24 hpi. The bar graph depicts the number of RAB7 spots <0.2 μm (blue) from the bacterial structure. Data show means ± SD, n=5 fields, 20-35 cells per field, from two independent experiments. Scale bar: 10 μm

### MIRO1, a Rho-family type GTPase, is required for enhanced MDV biogenesis in *p62^KD^* cells

Next, we wanted to understand what regulates MDV biogenesis in *p62^KD^*cells. Along with PRKN1 and DRP1, another family of proteins that have recently been implicated in MDV biogenesis is MIRO1/2 (50). At the same time, SNX9 is reported to help in the trafficking of MDVs (38). We knocked down DRP1 (DNML1), SNX9, MIRO1 (RHOT1), and MIRO2 (RHOT2) in separate experiments using WT and *p62^KD^* cells, and screened for conditions that could reverse the decline in *Mtb* CFU in the *p62^KD^* cells (Fig. S6A). Out of these, *siMIRO1* and *siMIRO2* were able to reverse the *Mtb* growth in the *p62^KD^* cells back to the levels seen in the WT cells (Fig. S6B-C, Fig. 7H). We next investigated whether the increase in the bacterial CFU in *MIRO1/2* knockdown cells was due to a direct role of MIRO1 and 2 in MDV biogenesis. We observed a significant decline in the cellular MDV count in *p62^KD^*cells where *MIRO1* or *MIRO2* were knocked down (Fig. 7I). This also led to a significant decline in the recruitment of TOM20 to *Mtb* phagosomes (Fig. 7J), which in turn led to reduced recruitment of RAB7 on *Mtb* phagosomes in the *MIRO1* or *MIRO*2 knocked down *p62^KD^* cells (Fig. 7K). Thus, MIRO1/2-mediated MDV biogenesis in *p62^KD^* cells contributes to enhanced TOM20 and RAB7 recruitment to *Mtb*-phagosomes.

## Discussion

Autophagy adaptors recognise and target the cargos for autophagy-mediated degradation (12). The damaged mitochondria are one of the significant cellular cargos that must get degraded via mitophagy to ensure mitochondrial quality and cellular health (51). Since the autophagy adaptors also play a crucial role in targeting the bacterial pathogens for lysosomal degradation, in this study, we wanted to study the distribution of the autophagy adaptors between mitochondria, the cellular cargo and *Mtb*, the pathogenic cargo, in the infected macrophages. Understanding this functional segregation of the autophagy adaptors is crucial since several host-directed therapies against tuberculosis envisage targeting the autophagy (52). However, most autophagy regulators do not distinguish between the homeostatic and defence arm of autophagy, potentially limiting the evolution of these regulators to clinically acceptable uses. Our data suggest p62/SQSTM1 as the major autophagy adaptor recruited to both *Mtb* and mitochondria in the infected macrophages. Contrary to our hypothesis, infection with *Mtb* did not significantly alter, except for a minor change in OPTN, recruitment and the distribution of autophagy adaptors on mitochondria. However, irrespective of their high or low association with the *Mtb* or mitochondria, individual depletion of each autophagy adaptor, except for NDP52, resulted in reduced bacterial survival in THP-1 macrophages. These early results showed that autophagy adaptors in the human macrophages regulate some housekeeping functions that crisscross with the anti-bacterial pathways. Moreover, it was also apparent that p62/SQSTM1 regulates some distinct downstream pathways in humans versus other vertebrates like mice and zebrafish. That is because, in the zebrafish model, p62/SQSTM1 and OPTN were previously shown to play a critical role in host resistance to mycobacterial infections (22). In mice, there are reports of an inverse correlation between *Mtb*-p62/SQSTM1 colocalisation and the bacterial CFU in the BMDMs (53). In yet another study, the absence of p62/SQSTM1 in mouse BMDMs did not affect Mtb CFU (54). The same study reported that starvation, which reduced Mtb CFU in the WT cells failed to do the same in the absence of p62/SQSTM1(54). In our experiments, mouse macrophages, knocking down autophagy adaptors in BMDM (*p62* and *Optn*) or knocking out in HOXB8-derived macrophages (p62), did not show any impact on intracellular *Mtb* growth, which is consistent with earlier reports (54). In addition, even *in vivo* studies in *p62^-/-^*animals did not show any significant impact on *Mtb* survival or killing (23). Curiously emerging evidences suggest that p62/SQSTM1could have tissue-specific functions (55). How p62/SQSTM1 regulates the anti-bacterial pathways in a host-specific manner has significant implications for developing new host-directed therapy against tuberculosis.

Autophagy, through mitophagy, ensures the maintenance of mitochondrial quality -a serious challenge for the cells with respiring mitochondria since the process of oxidative phosphorylation itself generates free radicals that can damage the mitochondria (17). The damaged mitochondria must get degraded via mitophagy to keep cellular apoptotic pathways under check (56). The damaged mitochondria can also cause bacterial killing, primarily through enhanced redox stresses (32, 57). While p62/SQSTM1 was highly recruited on mitochondria in the macrophages, we did not observe any loss to mitochondrial quality in *p62^KD^* cells. However, loss of p62/SQSTM1 in human cortical neurons is associated with an aberrant mitochondrial functionality (58). In HeLa cells, only OPTN, NDP52, and, to some extent, TAX1BP1 were essential for the mitophagy (59). Therefore, it is possible to have tissue/species-specific functions for each autophagy adaptor. Despite maintaining MQC, *Mtb* witnessed enhanced oxidative stress in the *p62^KD^*cells; which also contributed to the reduced bacterial survival in these cells. The increased redox stress could also be due to the proven role of p62/SQSTM1 in regulating the KEAP1-NRF2 antioxidant pathway (60, 61).

Interestingly, in addition to its role in mitophagy, p62/SQSTM1 also regulates peroxisome quality control by recognising ubiquitinated peroxisome protein PEX5 and targeting the peroxisomes for pexophagy (62). We cannot rule out peroxisomal dysfunction in *p62^KD^* cells, which could also contribute towards enhanced oxidative stress witnessed by the *Mtb*. The involvement of pexophagy in regulating redox stress during mycobacterial infections is only beginning to emerge and needs further explorations (63).

The more exciting discovery in this study was the determination that in the absence of p62/SQSTM1, these cells maintained mitochondrial quality through MDV biogenesis. As described by others, MDV biogenesis takes over in the absence of mitophagy to ensure MQC (37). The MDV biogenesis in *p62^KD^* cells was independent of infection, suggesting the homeostatic role of this process. Targeting MDVs to the lysosome for degradation ensures MQC and depends on RAB7 (64). Since MDV biogenesis in *p62^KD^* cells is independent of infection, we believe that RAB7 recruitment to *Mtb* phagosome occurs inadvertently during the transit of TOM20+-MDVs towards the lysosomes. Phagosome maturation arrest, or inhibition of xenophagy flux, caused by Mtb, specifically acts via the exclusion of RAB7 from the *Mtb*-containing phagosomes and autophagosomes (5) (45). Since RAB7 recruitment was the effector that impacted *Mtb* survival in *p62^KD^* cells, depletion of RAB7 rescued *Mtb* survival only in *p62^KD^* cells. Mechanistically, RAB7 depletion reduced the lysosomal targeting of *Mtb* and rescued *Mtb* from the redox stress, coinciding with a reduction in TOM20 recruitment to the *Mtb* phagosomes. Thus, MDV recruitment to *Mtb* phagosomes facilitated bacterial targeting to the lysosomes and also brought redox mediators to the bacterial compartments. At least one report in the literature suggests redox mediators brought in by MDVs to the *Staphylococcus*-containing phagosomes, which helps in the bacterial killing (46).

Could MDV biogenesis be blocked in *p62^KD^* cells to rescue *Mtb* survival? Among many regulators, Parkin, DRP1 and MIRO1/MIRO2 are implicated in MDV biogenesis (39, 50). Depleting Parkin in *p62^KD^* cells provided yet another intriguing insight. In the absence of Parkin, MQC was affected in both WT and *p62^KD^* cells, as expected from what is known about Parkin’s function in MQC (64). However, this also led to a further decline in bacterial survival in WT cells, contrary to earlier results using *PRKN^-/-^*BMDMs (65). Crucially, in the *p62^KD^*cells, the loss of Parkin synergised and further enhanced bacterial killing. Contrary to the perceived role of Parkin in MDV biogenesis, loss of Parkin in *p62^KD^* cells had no impact on TOM20 recruitment to *Mtb* phagosomes and lysosomes. In fact, increased *Mtb* killing in Parkin depleted condition was more consistent with other studies that show possible negative influence of Parkin on MDV formation (50, 66). In contrast, knocking down *MIRO1* and *MIRO2* in *p62^KD^*cells inhibited MDV biogenesis, reduced TOM20 and RAB7 recruitment on *Mtb* phagosomes and rescued intracellular *Mtb* CFU. It appears that cells can activate yet another MQC mechanism in *p62^KD^* cells independent of MIRO1/2. Further studies are needed to understand the complex interplay between diverse MQC pathways. Collectively, we conclude that the cumulative effect of lysosomal targeting and mitochondrial damage compromises bacterial survival in *p62^KD^* cells (Fig. 8).

**Fig. 8.**
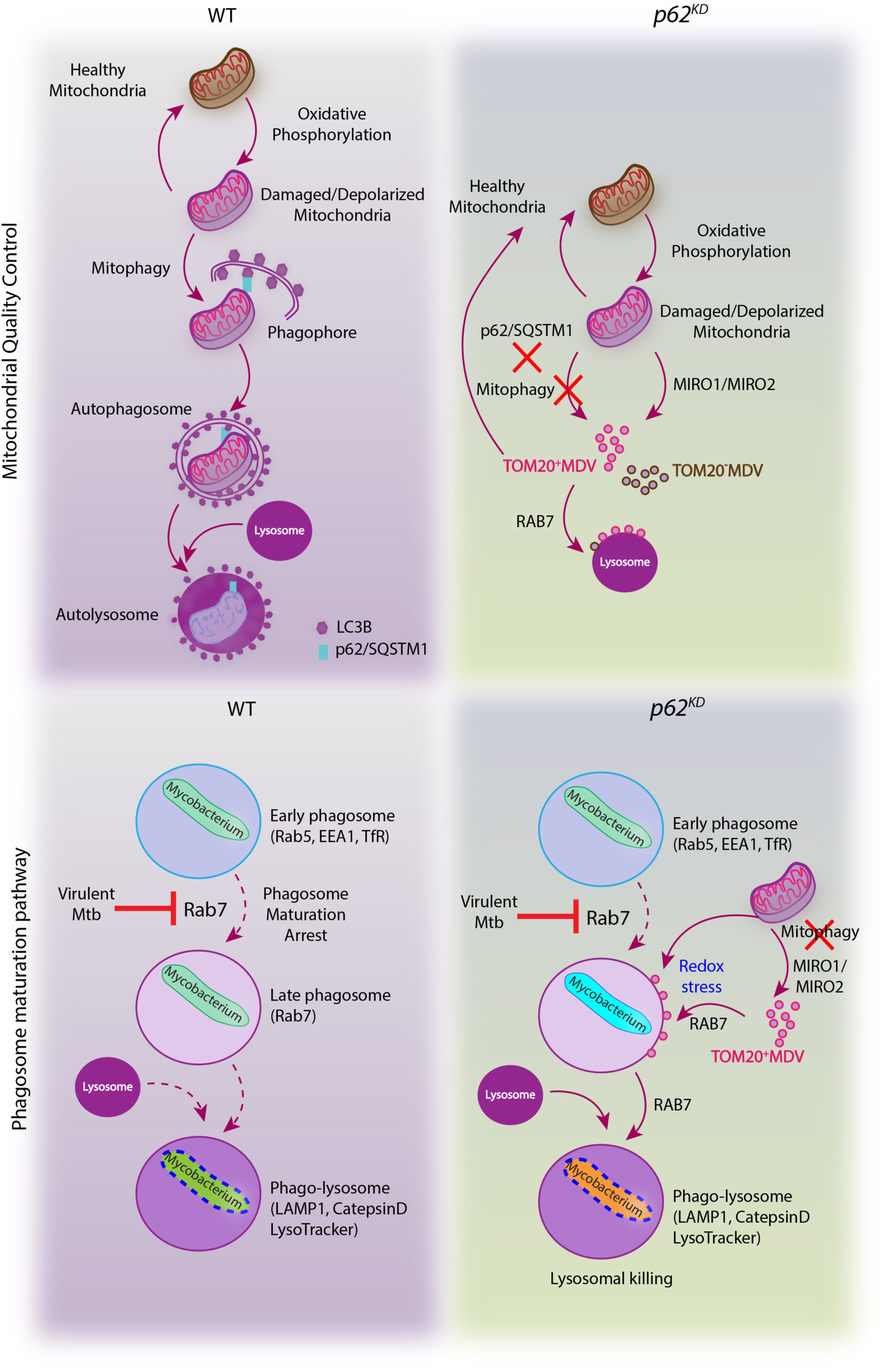
Crosstalk between mitochondrial quality control mechanisms and lysosomal targeting of *Mtb* in macrophages. Mitochondrial depolarisation triggers the mitophagy process by recruiting autophagy adaptor p62/SQSTM1 on mitochondria. p62/SQSTM1 binds the LC3B to form the mitophagosome, which fuses with lysosomes to clear the damaged mitochondria. **In the *p62^KD^* cells,** the loss of p62/SQSTM1 impairs the mitophagy process and triggers an alternative mitochondrial quality control pathway, i.e., the biogenesis of MDVs. MIRO1 and MIRO2 drive the formation of these vesicles from the outer mitochondrial membrane enriched with TOM20 (TOM20+-MDVs). They bud off from the damaged areas of mitochondria to fuse with lysosomes in a RAB7-dependent process. In WT macrophages, virulent strains of *Mtb* inhibit the RAB5 to RAB7 conversion, thereby preventing the fusion of early bacterial phagosomes/autophagosomes with the lysosomes, a process known as phagosome maturation arrest. In the *p62^KD^* condition, the TOM20^+^-MDVs get recruited on *Mtb* phagosomes/autophagosomes that inadvertently also help recruitment of RAB7 to the *Mtb*-phagosomes, thereby overcoming the phagosome maturation arrest. RAB7 recruitment to the phagosomes enhances the lysosomal targeting and *Mtb* killing in these cells. Both MDVs and damaged mitochondria contribute to the redox-mediated killing of *Mtb* and complement the lysosomal killing of *Mtb*.

This study provides an as-yet unrecognised crosstalk between mitochondrial quality control and the phagosome maturation pathway. The reversal of phagosome maturation reported here is unprecedented and underscores the need to dissect further the MQC pathways vis-à-vis anti-bacterial pathways in the host. We propose that these results provide novel ways to target MQC to achieve bacterial killing as a potential host-directed therapy strategy.

## Materials and Methods

### Ethics Statement

The Institutional Ethics Committee, ILBS (IEC/2020/81/MA03) and ICGEB (ICGEB/IEC/2020/25, version 3) approved studies on healthy human samples (healthy adults following informed consent). For BMDM experiments, animals were approved by IAEC of ICGEB (ICGEB/IAEC/18092021/CI-15).

### Cell Culture

Human monocytic cell line THP-1 was procured from American Type Culture Collection (ATCC) and cultured in RPMI 1640 medium (Cell Clone, CC3014) along with 10% Fetal Bovine Serum (Gibco). Cells were regularly checked for mycoplasma contamination. The cells were incubated in a humidified 5% CO_2_ chamber at 37 °C. THP-1-derived macrophages were obtained by treating THP-1 cells with 20 ng/ml phorbol myristate acetate (Sigma, P8139) for 24 hours, followed by washing and maintenance in complete media. Human MDMs (hMDMs) were isolated from human PBMCs and separated from healthy human donors’ blood using PolymorphPrep (PROGEN,1895) layering and centrifugation. The heparinised blood was layered on PolymorphPrep in a 1:1 ratio and centrifuged at 500g for 45 min at 25°C. The interface containing PBMC was isolated carefully and treated with RBC lysis buffer. Subsequently, cells were washed twice with DPBS. PBMCs were diluted in 10% RPMI 1640 media to a concentration of 1 × 10^6^ cells/ml. Cells were put in a six-well cell culture plate and incubated for 4 hours at 37°C in a humidified 5% CO_2_ chamber. Non-adherent cells were removed, followed by two washes with RPMI media. Complete RPMI media containing 50 ng/ml recombinant human M-CSF (R & D systems, 216-MC-025) was added and cells were allowed to differentiate for a week into macrophages in a humidified 5% CO_2_ incubator at 37 °C. For BMDMs, bone marrow was isolated from the femur and tibia (hind limb) of mice (6-8 wk old, female, C57BL/6 mice) and passed through a 23G needle. The RBCs were lyzed by RBC lysis buffer and then washed with 1x PBS. The isolated Bone marrow cells were cultured in 10% DMEM containing 20% L929 supernatant for 5-7 days. All mice were housed in the Bio experimentation animal facility in a sterile environment.

### Generation of Hoxb8 progenitors cell lines and differentiation in macrophages

The method was previously described (67). Briefly, BM cells were collected from the femurs of 8-week-old mice and separated on a Ficoll gradient. Cells were resuspended in RPMI containing 15% FBS and supplemented with 15% of Flt3L and 30% GM-CSF containing supernatant. After 2 days of cell culture, cells were collected and resuspended in RPMI supplemented with 10% FBS, 1 μM β-estradiol (MilliporeSigma, E-2758), 1% gentamicin and infected with MSCV herad-3HAHOXB8 retroviral particles. For 1-month, cells were dispensed in fresh medium every 3–4 days and transferred into new wells. After this period, cells were expanded in Hoxb8 Medium supplemented with 15% Flt3L supernatant. For subsequent differentiation experiments, cells were washed twice with warm PBS containing 1% FBS and resuspended in a concentration of 0.5 × 10^5^ cells/mL in IMDM containing 20% M-CSF. On day 4 medium was replaced with a fresh one, and 1/3 of starting media was added on day 6. Differentiated cells were used between days 7 and 8. Flow cytometry confirmed homogeneous differentiation between different genotypes using F4/80 and CD11b markers.

#### Mycobacterium tuberculosis culture and infection

Mycobacterial strain *H37Rv* was obtained from Colorado State University. The mCherry-*H37Rv* was made using pMSP12: mCherry plasmid (Addgene No. 30169). *H37Rv* culture was grown in Middlebrook 7H9 (BD, 271310) media supplemented with 10% ADC (Albumin Dextrose Catalase, BD, 211887), 0.4% glycerol, and 0.05% Tween 80 until the mid-log phase.

The bacteria were then pelleted down by spinning at 4000 rpm for 5 min, washed with RPMI 1640 and their single-cell suspension was obtained by passing the culture through needles of 3 different gauge (G) sizes in the following way- 23 G (7 times), 26 G (5 times) and 30 G (3 times). For *Mtb* quantification, OD600 was considered equivalent to 100 million bacteria. The required number of bacteria was added onto the seeded cells in complete media at MOI of 1:10, followed by spinning the plates at 800 rpm for 5 min. After 4 hrs, 200ug/ml amikacin sulphate (Sigma, A2324) treatment was done for 2 hrs. Subsequently, cells were washed, fresh media was added, and cells were kept at 37°C for the desired time point of processing. Thus, the duration to complete the infection protocol was 6 hours, and the time point immediately after the amikacin step was considered 0 hours post-infection (hpi). All *Mtb* work was carried out in the Tuberculosis Aerosol Challenge Facility (TACF), a national BSL3 facility for tuberculosis research.

### CFU Experiment

For CFU experiments, the cells were seeded in 96-well cell culture plates. At desired time point, culture media was removed, and cells were washed with 1X PBS. Cells were lysed with 100 μl of 0.06% SDS lysis buffer for 10 minutes at room temperature. The lysate was then diluted in 7H9 media, and 10μl of this sample was plated on 7H11 agar (BD, 8801671) square plates supplemented with 10 % OADC (BD, 211886). Plates were dried and incubated at 37°C humidified incubator for a minimum of two weeks or till the appearance of *Mtb* colonies. CFUs were calculated by multiplying the actual colony count with the dilution factor and the volume of the diluted sample used for plating. For the replication clock experiment, pBP10 containing H37Rv was used for infection, and at the designated time points, CFU plating was performed parallelly on normal or kanamycin-containing 7H11-OADC plates. The fraction of CFU obtained in Kan^+^ plates compared to the Kan^-^ plates was calculated to determine the %pBP10^+^ *Mtb*.

### Flow Cytometry Assays

Cells were seeded in a 12-well culture plate for all the flow cytometry experiments. The H37Rv used was labelled with CellVue Claret Far red dye (Sigma, MINCLARET-1KT) according to the manufacturer’s protocol. After processing the samples at the desired time, 500 μl of 1X PBS was added to the wells, and cells were scrapped off using a scrapper and added to fresh round bottom polystyrene FACS tubes. The samples were immediately acquired in BD FACSAria Flow cytometer. For most experiments, 30000 events were captured and analysed by FlowJo v10 software. The assay for DCFDA (10 μM, 30 minutes) (Sigma, D6883), TMRE (100 μM, 30 minutes) (Invitrogen, T669), Mitotracker Green (200 nM, 40 minutes) (Invitrogen, M7514), and MitoSOX Red (5 μM, 20 minutes) (Invitrogen, M36008) were performed according to manufacturer’s protocol.

### Immunoblotting

For the lysate preparation, cells were seeded in a 6-well cell culture plate, and at the desired time, cells were washed with ice-cold 1X PBS. The lysis of the cells was performed by incubating them with the lysis solution containing RIPA buffer (Sigma-Aldrich, R0278) and 1X Protease Arrest (G^−^Biosciences) for 20 minutes on ice. The lysate was collected in microcentrifuge tubes and centrifuged at 6000 *g* for 10 minutes at 4 °C. The supernatant (cytoplasmic extract) was collected, and protein quantification was done by Bradford Assay. The protein sample was mixed with 6× loading dye, and denaturation was done at 95 °C for 10 minutes. The protein sample was subjected to SDS-PAGE for separation and blotted onto the nitrocellulose membrane (BIO-RAD) using BIO-RAD Trans-Blot Turbo Transfer System.

Blocking was performed for an hour with Odyssey blocking buffer (LI-COR Biosciences) and 1X PBS in the ratio of 1:1 at room temperature. Blots were stained overnight with primary antibody at 4 °C and then with secondary antibody for 1.5 hours at RT, with good washes with 1X PBST in between. The IR-conjugated secondary antibodies anti-mouse IR780 (926–32210) and anti-rabbit IR780 (925–68071) were obtained from LI-COR Biosciences. The dilutions of the antibodies were made according to the supplier’s protocol in the Odyssey blocking buffer. The blots were scanned with the Odyssey Infra-Red Imaging system (LI-COR Biosciences).

### siRNA Transfection

Cells were either transfected with control or specific siRNA using Dharmafect-2 reagent (Dharmacon Inc, T-2002-02) according to the manufacturer’s protocol. 50nM siRNA was added to cells at 0-hour post infection for 24 hours. All the siRNA used in our study were siGenome siRNA SmartPool (Dharmacon Inc). The following were used: p62/SQSTM1 (M-010230-00-0005), NDP52 (M-010637-01-0005), OPTN (M-016269-02-0005), TAX1BP1 (M-016892-01-0005), NBR1 (M-010522-01-0005), RAB7A (M-010388-00-0005), PARKIN (PRKN) (M-003603-00-0005), MIRO1 (RHOT1) (M-010365-01-0005), MIRO2 (RHOT2) (M-008340-01-0005), DRP1 (DNM1L) (M-012092-01-0005), and SNX9 (M-017335-02-0005). In BMDM, mouse-specific p62/sqstm1 (M-047628-01-0005) and optn (M-041705-01-0005) siRNA was used.

### Confocal Microscopy

For microscopy experiments, cells were seeded on the sterile coverslips in a 24-well tissue culture plate. All the staining steps were performed at room temperature. At the requisite time, cells were fixed with 4% PFA (Sigma, 158127) for 20 minutes, followed by 2 washes with 1X PBS. The cells were permeabilised with 0.2% Triton-X100 (Sigma, T8787) (in 1X PBS) for 15 minutes with two subsequent washes with 1X PBS. Blocking was performed with the buffer of 3% BSA (Sigma, A7906) and 0.5% Tween 20 (Sigma, P1379) in 1X PBS for 1 hour. The cells were stained with respective primary antibodies for 2 hours and then washed twice with 1X PBST. This was followed by staining with individual secondary antibodies tagged with the fluorophore of choice for 1 hour. Both primary and secondary antibodies were made in the blocking buffer at the required dilution. The cells were washed twice with 1X PBS before mounting the coverslips on glass slides with ProLong Gold anti-fade reagent (Life Technologies, P36930). Confocal images were acquired with a Nikon A1R microscope equipped with 100X/1.4 NA plan-apochromat VC, DIC N2 objective lens. The analysis was performed on Imaris 9.6 imaging software using several tools like deconvolution, 3D reconstruction and shortest distance calculation. The research for the roGFP2 redox sensor was conducted with Nikon NIS Element. For MDVs spot analysis, Deep learning-based TruAI tool of cellSens Imaging software was used.

The primary antibodies used were: α-p62/SQSTM1 (Abcam, ab155686), α-OPTN (Abcam, ab23666), α-NDP52 (Abcam, ab68588), α-TAX1BP1 (Abcam, ab121812), α-NBR1 (Novus Biologicals, NBP1-71703), α-MAP1LC3B (NB100-2200), α-BNIP3 (Abcam, ab109362), α-FUNDC1 (ab74834), α-NIX (Abcam, ab8399), α-TOM20 (Abcam, ab56783, ab78547), α-Parkin (Novus Biologicals, NBP2-67017), α-HSP60 (Abcam, ab13532), α-SOD2 (Novus Biologicals, NB100-1992), α-PDHA1 (Abcam, ab110330), α-DRP1 (Abcam, ab184247), α-SNX9 (Abcam, ab181856), α-ECSIT (Novus Biologicals, NBP1-918858), α-LAMP1 (Santa Cruz Biotechnology, Sc-20011), MIRO1 (Novus, NBP1-89011), MIRO-2 (Novus, NBP1-88982), α-RAB7 (Abcam, ab50533), α-CATHEPSIN-D (Santa Cruz Biotechnology, Sc-10725), α-Ubiquitin (Abcam, ab134953), α-MCU (Cell Signaling Technology, D2Z3B), α-TUBULIN (Abcam, ab6046), α-ACTIN (Abcam, ab8226), α-GAPDH (Abcam, ab9485)

All secondary antibodies used were from Invitrogen, conjugated with Alexa Flour dye: anti-mouse Alexa Fluor 405 (A31553), anti-mouse Alexa Fluor 488 (A28175), anti-mouse Alexa Fluor 568 (A11031), anti-mouse Alexa Fluor 647 (A21235), anti-rabbit Alexa Fluor 405 (A31556), anti-rabbit Alexa Fluor 488 (A11034), anti-rabbit Alexa Fluor 568 (A11011), anti-rabbit Alexa Fluor 647 (A21245). CellROX deep red reagent (C10422) and LysoTracker green (L7526) were from Invitrogen. For microscopy experiments, mCherry-*H37Rv* was used.

### *p62^KD^* THP-1 monocytic cell line generation

The *p62^KD^* cells were generated using the CRISPR/Cas9 system. The guide RNA was designed against exon 3 of p62 and cloned into a gRNA cloning vector plasmid (Addgene No. 41824). The Cas9 enzyme encoding hCas9 plasmid was from Addgene (No. 41815). Both the plasmids were co-transfected into THP-1 monocytes using the Invitrogen Neon Transfection system according to the manufacturer’s protocol. At 24 hours post-transfection, the cells were selected on Neomycin (Sigma, G418 A1720) at 750ug/ml for 21 days, followed by checking the absence of p62 in knockdown cells by microscopy and immunoblotting.

### RNA Sequencing

Total RNA was extracted from THP-1 macrophages, following the double extraction protocol: RNA isolation by acid guanidinium thiocyanate-phenol-chloroform extraction (TRIzol) followed by a Qiagen RNeasy Micro clean-up procedure. RNA was analysed on Agilent Bioanalyser for quality assessment with RNA Integrity Number (RIN) ranging from 7.5 to 10 and a median of RIN 9.8. cDNA libraries were prepared using 2 ng of total RNA using the SMARTSeq v2 protocol with the following modifications: 1. Addition of 20 µM TSO; 2. Use of 200 pg cDNA with 1/5 reaction of Illumina Nextera XT kit. The length distribution of the cDNA libraries was monitored using a DNA High Sensitivity Reagent Kit on the Perkin Elmer Labchip. All samples were subjected to an indexed paired-end sequencing run of 2x151 cycles on an Illumina HiSeq 4000 system (28 samples/lane).

### RNA-Seq analysis

Reads were quality filtered using Trimmomatic (v.0.39, LEADING:10 TRAILING:10 SLIDING WINDOW:4:15 MINLEN:75). They were assessed for quality using FastQC (v.10.1))] and visualised with MultiQC (v.1.9). Salmon (v.1.8, –numGibbsSamples 30) was used to quantify quality filtered reads against the human genome (GRCh38, primary assembly) with Gencode annotation (v.40). Profiles were imported using tximeta (v.1.10) and SummarizedExperiment (v.1.22) which were then summarised to gene-level expression. Non-coding genes were filtered out, and the following gene biotypes were retained: protein-coding, pseudogenes (translated and transcribed), Immunoglobulin (IG) and T cell receptor (TCR) genes). Dataset was further filtered to retain only those genes whose expression was detected in at least 3 samples. Statistical analysis was done using swish methodology from fishpond (v.1.8) (68) and significantly differentially expressed genes (DEGs) were defined based on *qvalue* ≤ 0.1 and absolute log_2_ fold change ≥ 1. Heatmap of the DEGs was generated with heatmap (v.1.0.12) using mean z-scaled normalised scaled TPM values (summarised to gene level using summarizeToGene() function) and clustered based on Euclidean distance and Ward.D2 methodology. Enrichment analysis was done separately on overexpressed and repressed DEGs from each pairwise comparison using enrichR (v.3.0) (69) with KEGG Human 2021 and Gene Ontology (GO) (70) (Biological Process (BP), Cellular Component (CC) and Molecular Function (MF)) 2021 database. Mitochondrial DEGs and their associated pathways were identified using Mitocarta 3.0 (71). Cytokines were identified from the comprehensive list as described by Pro *et al.*(*72*). Figures were generated using ggplot2 (v.3.3.5). Analyses were done in R statistical software (v.4.1).

### Mrx1-roGFP2 Redox Sensor Assay

*H37Rv* cultures were electroporated with Mrx1-roGFP2, and colonies were selected on a hygromycin 7H11 plate.THP-1 macrophages were infected with Mrx1-roGFP2-*H37Rv* at M.O.I of 10. At the requisite time, cells were washed with 1X PBS, with subsequent treatment of 10 mM NEM (N-ethylmaleimide, Sigma, E3876) for 10 minutes at 37 °C. To the positive and negative control groups, 5 mM CHP (Cumene hydroperoxide, Sigma) and 10 mM DTT (Dithiothreitol, Sigma, D0632) were added respectively for 5 minutes at 37 °C. The cells were washed twice with 1X PBS, followed by NEM-PFA fixing. For FACS, 2% PFA was used, whereas for confocal microscope acquisition, 4% PFA was used.

### Extracellular flux analysis

At 24 hours post-infection, THP-1 macrophages were scrapped off from a 6-well culture plate, and 4 x 10^4^ cells were seeded in Cell-Tak (Corning, 354240) coated Agilent Seahorse XFp cell culture mini plate (Agilent, Santa Clara, CA). The plates were centrifuged at 200g for 1 min with minimum deceleration. The plates were parafilm-wrapped and incubated in a non-CO_2_ incubator at 37°C for 25-30 minutes. After adding 130μl warm assay media and incubation for for 15-25 minutes at 37°C, the plates were ready for the assay. The Agilent Seahorse XFp analyser was used to assess the oxygen consumption rate. The modulators included in this assay were oligomycin (1.5 μM, Sigma, #75351), Carbonyl cyanide-4 (trifluoromethoxy) phenylhydrazone (FCCP) (1 μM; Sigma, C2920), Rotenone (0.5 μM; Sigma, R8875) and antimycin A (0.5 μM; Sigma, A8674) (73). Initially, the baseline OCR of the cells was measured, from which the basal respiration was determined by eliminating the non-mitochondrial respiration. Maximal respiration was calculated by subtracting the non-mitochondrial respiration from the maximal OCR after FCCP addition. Non-mitochondrial respiration represents the minimum OCR after rotenone and antimycin A treatment.

### mtDNA/nDNA

The cellular DNA was extracted from the DNeasy kit (Qiagen) according to the manufacturer’s protocol. To quantify the mtDNA/nDNA ratio, qPCR was performed to amplify one gene from the mitochondrial genome (mt*CO1*) and one gene from the nuclear genome (*RPL13A*). Primer sequences were as: mt*CO1*_forward: 5′-CAGGAGTAGGAGAGAGGGAGGTAAG-3′, mt*CO1_*reverse: 5′-TACCCATCATAATCGGAGGCTTTGG-3′; *RPL13A*_forward:5′-CGCCCTACGACAAGAAAAAG-3′, and *RPL13A*_reverse: 5′ CCGTAGCCTCATGAGCTGTT-3′

### Inhibitor or Activators treatment

The reagents were used according to the following conditions before processing the samples: Bafilomycin A1 (B1793) 100 nM for 3 hrs, Carbonyl Cyanide m-Chlorophenylhydrazine (CCCP, Sigma, C2759): 5 μM for 12 hours for CFU and 20 μM for 20 min for mitochondrial depolarisation assay (TMRE), Nigericin (Sigma, N7143-5MG): 2 μM for 24 hours, N-acetyl cysteine (NAC, Sigma, A7250), 2mM for 24 hours.

### RT-PCR

Total RNA was extracted at the required time point from THP-1 macrophages using MDI total RNA Miniprep kit (MTRK250) according to the manufacturer’s protocol. cDNA was synthesised from 500 ng of total RNA by reverse transcriptase PCR using a BIO-RAD iScript cDNA synthesis kit (170–8891). The qPCR was set using the mixture of cDNA sample, SYBR green dye (QuantiNova) and desired primers, where 18s rRNA was used as an internal control. The acquisition was made on the Bio-Rad CFX 96 Real-time PCR system. The primers used are *Fw_p62/SQSTM1: 5’*-*GCACCCCAATGTGATCTGC-3’*,

*Rv_p62/SQSTM1:5’-CGCTACACAAGTCGTAGTCTGG-3’,*

*Fw_NBR1:5’-AGGAGCAAAACGACTAGCTGC-3’,*

*Rv_NBR1:5’-TCTGGGGTCTTCATGTCTGAT-3’,*

*Fw_NDP52:5’-ATTTCATCCCTCGTCGAAAGGA-3’,*

*Rv_NDP52:5’-TGAAGGTGTAATACTCACGGGTT-3’,*

*Fw_OPTN:5’-CCAAACCTGGACACGTTTACC-3’,*

*Rv_OPTN:5’-CCTCAAATCTCCCTTTCATGGC-3’,*

*Fw_TAX1BP1:5’-TAAAGGAGCAACTTCGTAAAGCA-3’,*

*Rv_TAX1BP1:5’-CTGCCATCGTTCTGTCTCGT-3’,*

*Fw_MFN1:5’-TGGCTAAGAAGGCGATTACTGC-3’,*

*Rv_MFN1:5’-TCTCCGAGATAGCACCTCACC-3’,*

*Fw_MFN2:5’-CTCTCGATGCAACTCTATCGTC-3’,*

*Rv_MFN2:5’-TCCTGTACGTGTCTTCAAGGAA-3’,*

*Fw_DRP1:5’-CTGCCTCAAATCGTCGTAGTG-3’,*

*Rv_DRP1:5’-GAGGTCTCCGGGTGACAATTC-3’,*

*Fw_OPA1:5’-TGTGAGGTCTGCCAGTCTTTA-3’,*

*Rv_OPA1:5’-TGTCCTTAATTGGGGTCGTTG-3’,*

*Fw_FIS1:5’-GTCCAAGAGCACGCAGTTTG-3’,*

*Rv_FIS1:5’-ATGCCTTTACGGATGTCATCATT-3’,*

*Fw_18s:5’-GCTTAATTTGACTCAACACGGGA-3’,*

*Rv_18s: 5’-AGCTATCAATCTGTCAATCCTGTC-3’*.

### Statistical Analysis

Statistical significance for comparisons between two sets of the experiment was performed using an unpaired two-tailed Student’s *t-test*. For multiple treatment experiments, one-way ANOVA followed by multiple comparison analyses was conducted in GraphPad PRISM 8.

## Data availability statement

The RNAseq data presented in this study is available at GEO database under the accession ID: GSE216985.

## Acknowledgements

We acknowledge Dr David Sherman for the replication clock plasmid (pBP10). We acknowledge the Tuberculosis Aerosol Challenge Facility (TACF), supported by the Department of Biotechnology, Govt of India (BT/INF/22/SP43067/2021), for all BSL3 experiments. This work was supported by the Department of Science and Technology through a DST-UKIERI collaborative grant (DST/INT/UKP-136/2016, DK and SS) and partly by the Department of Biotechnology, Govt of India (BT/IC-06/003/91 and RAD-40/104/2023-MED-DBT; Flagship program, DK), DBT-Wellcome Trust India Alliance Senior Fellowship (IA/S/17/1/503071, DK) and Science and Engineering Research Board (SERB), Govt of India (EMR/2016/005296, and CRG/2022/008256; DK). University of Birmingham India Institute Visiting Fellowship to S.V. and S.S.; LifeArc, Wellcome Trust and Birmingham Fellowship to S.S. and DST-INSPIRE fellowship to S.V. A.S is supported by A*STAR ID Labs, R01HL152078 and NMRC/OFIRG/may-0096.

## Author contributions

Conceptualization: DK, SS, SV

Methodology: SV, ST, RS, VY, GMP, MZK, MM, SF, SH, PS, ST

Investigation: SV, ST, RS, MD, VY

Visualization: SV, ST, MD, DK

Resources: VN, BM, MB, JSM, FB

Supervision: DK

Writing—original draft: DK, SV

Writing—review & editing: DK, SV, ST, MD, VY, FB, AS, SS, AS

## Competing interests

All authors declare they have no competing interests.

## Data and materials availability

All data are available in the main text or the supplementary materials. The RNAseq raw data is available at GEO database, accession number- GSE216985

## Supporting information

**S1 Fig. Expression profile of autophagy adaptors in H37Rv infected and uninfected THP-1 Macrophages.** (A) Gene expression of NBR1, NDP52, TAX1BP1, OPTN, and p62/SQSTM1 (p62) by RT-PCR at 0-, 24-, and 48-hpi. Data show mean ± SD from three independent experiments. (B) Immunoblots represent protein expression of TAX1BP1, NDP52, OPTN, NBR1, and p62 at 0-,24- and 48-hpi; GAPDH is used separately as the loading control for each condition; (C) bar graphs on the right show the quantification of the protein levels of these adaptors after normalising with GAPDH. Data show mean ± SD from three independent experiments.

**S2 Fig. Depletion of autophagy adaptors in human and mouse macrophages and their effect on H37Rv survival.** (A) Immunoblot represent the siRNA-targeted depletion of adaptors TAX1BP1, p62, NDP52, OPTN and NBR1 in THP-1 monocytes in the separate experiment for each adaptor at 24 hours post-transfection. The numbers depict the ratio of intensities of specific proteins to that of GAPDH. (B) CFU analysis of H37Rv in control and siRNA-targeted-OPTN and -p62 U937 macrophages at 24 hpi. Right panel: Immunoblot represents the siRNA-targeted depletion of p62 and OPTN at 24 hours post-transfection. The numbers depict the ratio of intensities of specific proteins to that of GAPDH. (C-D) Respective immunoblot and IFA images for p62 in *p62^KD^* THP-1 cells generated using CRISPR/Cas9 and their quantification graphs (Right Panel). Plots represent data from three experiments (C) and 5 fields from two different experiments (D). (E) Immunoblot depicts the levels of p62 in uninfected and infected cells in WT and *p62^KD^* cells at 24- and 48 hpi and separate quantification plots from two independent experiments (Right). (F) Immunoblot represents LC3B levels in the lysates of WT and *p62^KD^* macrophages at 0-,24-, and 48-hpi in the presence and absence of BafA1 (100nM for 3 hours before processing). (G) Confocal images show the interaction of LC3B spots (blue spots, at <0.2 um from bacteria) and H37Rv. The bar graph represents the quantification of blue spots on H37Rv in WT and *p62^KD^*cells in the presence and absence of BafA1 at 0- and 24-hpi. Data show mean ± SD, n> 9 Fields, 20-35 per field, from three experiments. (H-I) Respective CFU analysis of H37Rv in control and siRNA-targeted p62/sqstm1 depleted and Optn depleted mouse bone marrow monocyte-derived macrophages at 24 hpi. Data show mean ± SD, from three independent experiments, right panel shows immunoblot confirming the p62 and OPTN depletion upon siRNA transfection. The numbers depict the ratio of intensities of specific proteins to that of GAPDH. (J) CFU analysis of H37Rv in WT and *p62^KO^*HOXB8 derived macrophages at 24 hpi. Data show mean ± SD, from three independent experiments. The immunoblot shows the levels of p62 in these cells.

**S3 Fig. Gene expression patterns and functional class enrichment.** (A) DEGs (differentially expressed genes, absolute log2FC ≥1 & q-value ≤ 0.1) at each time point. Row annotation represents DEGs identified when compared between infected versus uninfected in WT and *p62^KD^* macrophages at 0-, 24- and 48-.hpi. Row clustering was done using ward. D2 methodology on Euclidean distance calculated from the z-scaled values. Heatmap key reflects expression intensity based on z-scaled values (B) shows Gene Ontology Biological Process (GOBP) enrichment of the upregulated or downregulated genes identified based on infected versus uninfected in WT and *p62^KD^ m*acrophages at 0-, 24- and 48-.hpi. Significance is determined by nominal p.value ≤ 0.05. Only the top 15 enriched GOBPs from each comparison are visualised. Coverage represents the proportion of genes identified in a particular GOBP out of the total genes associated with that process. Colour intensity represents the nominal p-value gradient. (C) is a subset of (A) which shows an expression of the identified DE cytokine genes between infected versus uninfected in WT and *p62^KD^* macrophages at 0-, 24- and 48-.hpi.

**S4 Fig. Impact of p62/SQSTM1 depletion on mitochondrial DNA content and fission-fusion pathways.** (A) Quantification of mtDNA by normalising the gene expression of the mitochondria encoded protein (mtCO1) with nuclear genome encoded RNA polymerase (RPL13A) in WT and *p62^KD^*THP-1 macrophages in infected and uninfected THP-1 macrophages at 0-, 24-, and 48-hpi. (B) Fold change in gene expression of mitochondrial fusion proteins MFN1, MFN2 and OPA1, and fission proteins DRP1 and FIS1 by RT-PCR in H37Rv infected WT and *p62^KD^* THP-1 macrophages at 24 hpi. (C) Immunoblots represent protein expression of BNIP3 in infected WT and *p62^KD^* cells at 24 hpi.

**S5 Fig. Effect of mitochondrial quality control in H37Rv infected THP-1 macrophages.** (A-E) Confocal images depict the interaction of TOM20 (green) with proteins from three different mitochondrial compartments (red). Representative TOM20 3D structures and mitochondrial protein spots: (A) PDHA1, (B) HSP60, (C) SO2, (D) ECSIT, and (E) MCU are shown at the right. The yellow spots are at <0.2 μm, and the magenta spots are at > 0.2 μm from TOM20 structures. (F) Immunoblot show the levels of RAB7 in siControl and siRAB7 in THP-1 macrophages at 24 hours post-siRNA transfection. (G) Immunoblot show the levels of PARKIN in siControl and siPRKN in THP-1 macrophages at 24 hours post-siRNA transfection. (H) Per cent MFI change in TMRE of infected THP-1 macrophages in the presence and absence of nigericin. (I) The relative oxidation state of Mrx1-roGFP2 reporter as the ratios of emission MFI at 510nm when excited at 405nm to 488nm in nigericin-treated WT and p62KD cells at 24 hpi. Data show mean ± SD, n=3. (J) H37Rv CFU in Nigericin-treated and untreated WT and p62KD cells. Data show mean ± SD, n=4, from two independent experiments. (K) The graph represents the MTT cell viability assay of nigericin-treated and untreated WT and p62KD THP-1 macrophages. Scale bar: 10 μm.

**S6 Fig. Role of MIRO1 and MIRO2 in MDV biogenesis.** (A) CFU analysis of H37Rv in control and siRNA-targeted-MIRO1, MIRO2, DNM1L (DRP1) and SNX9 WT and *p62^KD^* macrophages at 24 hours post-transfection, Data show mean ± SD, n=5, from two independent experiments. (B-C) Immunoblot represents the siRNA targeted depletion of MIRO1 and MIRO2, respectively at 24 hours post-transfection.

**S1 Table. Infection-associated changes at different hours**

**S2 Table. Enrichment analysis of the DEGs**

**S3 Table. DEGs identified to be associated with mitochondria based on Mitocarta 3.0 (a subset of total DEGs)**

**S4 Table: Mitochondrial DEGs associated with mitochondria pathways based on Mitocarta 3.0**

**S5 Table: DEGs identified to cytokine genes (a subset of total DEGs)**

## Notes

### Competing Interest Statement

The authors have declared no competing interest.

### Summary of Updates

The main change in the current version, compared to the previous version, is the results in Fig. 2C. The replication clock plasmid data is presented in way that allows better interpretation of differences seen between the WT and p62KD cells.

