## Supplementary material for "A mitochondrial quality control mechanism reverses the phagosome maturation arrest caused by *Mycobacterium tuberculosis*": Figure S1-S6

Surbhi Verma *et al.*

**This PDF file includes:**

Figs. S1 to S6
Title of Table S1 to S5

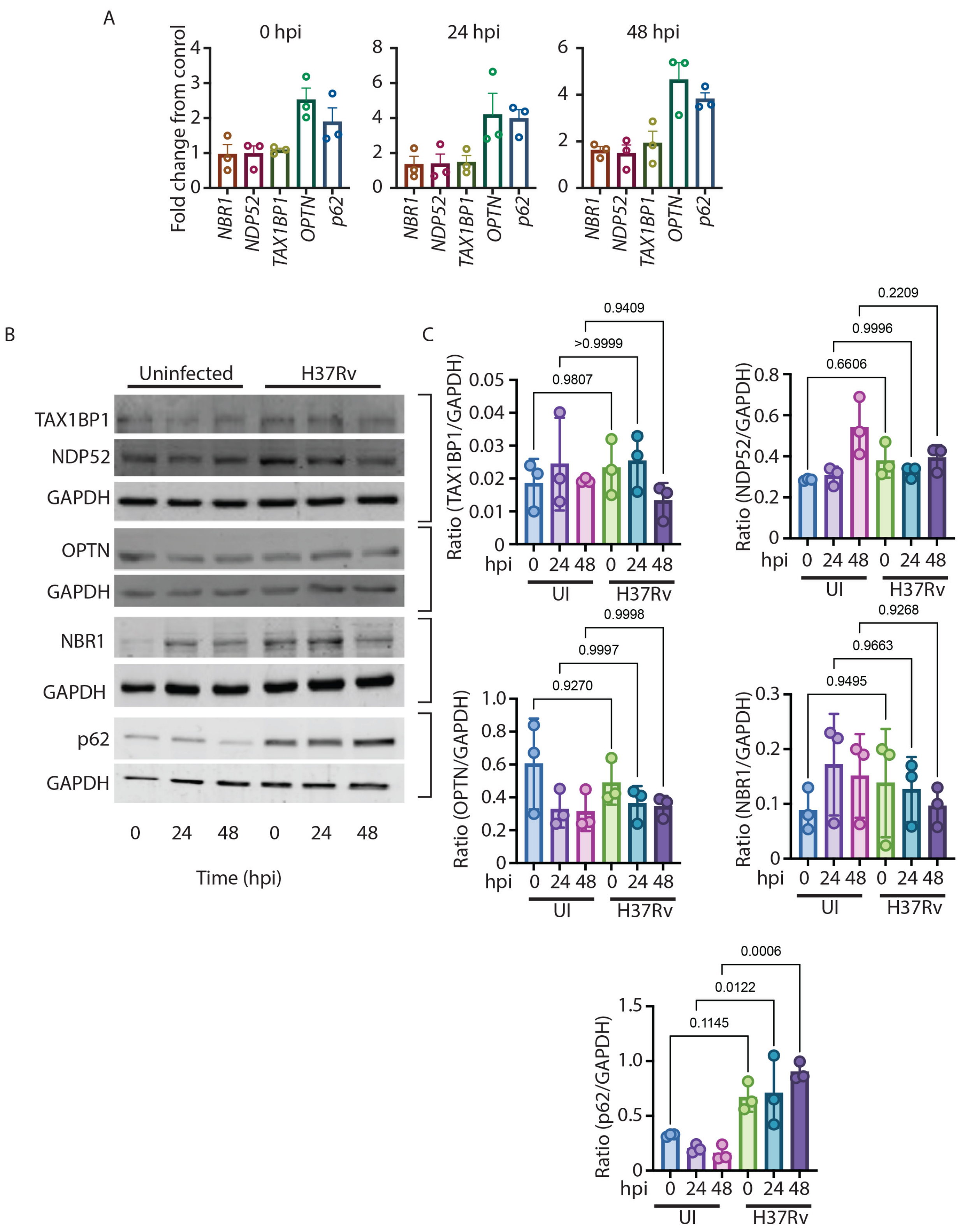
A
0 hpi
24 hpi
48 hpi
Fold change from conrol
NBR1
NDP52
TAX1BP1
OPTN
p62
B
Uninfected
H37Rv
TAX1BP1
NDP52
GAPDH
OPTN
GAPDH
NBR1
GAPDH
p62
GAPDH
0 24 48 0 24 48
Time (hpi)
C
Ratio (TAX1BP1/GAPDH)
0.9409
>0.9999
0.9807
Ratio (NDP52/GAPDH)
0.2209
0.9996
0.6606
Ratio (OPTN/GAPDH)
0.9998
0.9997
0.9270
Ratio (NBR1/GAPDH)
0.9268
0.9663
0.9495
Ratio (p62/GAPDH)
0.0006
0.0122
0.1145
hpi 0 24 48 0 24 48
UI
H37Rv

**Fig. S1.**

**Expression profile of autophagy adaptors in H37Rv infected and uninfected THP-1 macrophages**

**(A)** Gene expression of NBR1, NDP52, TAX1BP1, OPTN, and p62/SQSTM1 (p62) by RT-PCR at 0-, 24-, and 48-hpi. Data show mean ± SD from three independent experiments. **(B)** Immunoblots represent protein expression of TAX1BP1, NDP52, OPTN, NBR1, and p62 at 0-, 24- and 48-hpi; GAPDH is used separately as the loading control for each condition; **(C)** bar graphs on the right show the quantification of the protein levels of these adaptors after normalising with GAPDH. Data show mean ± SD from three independent experiments.

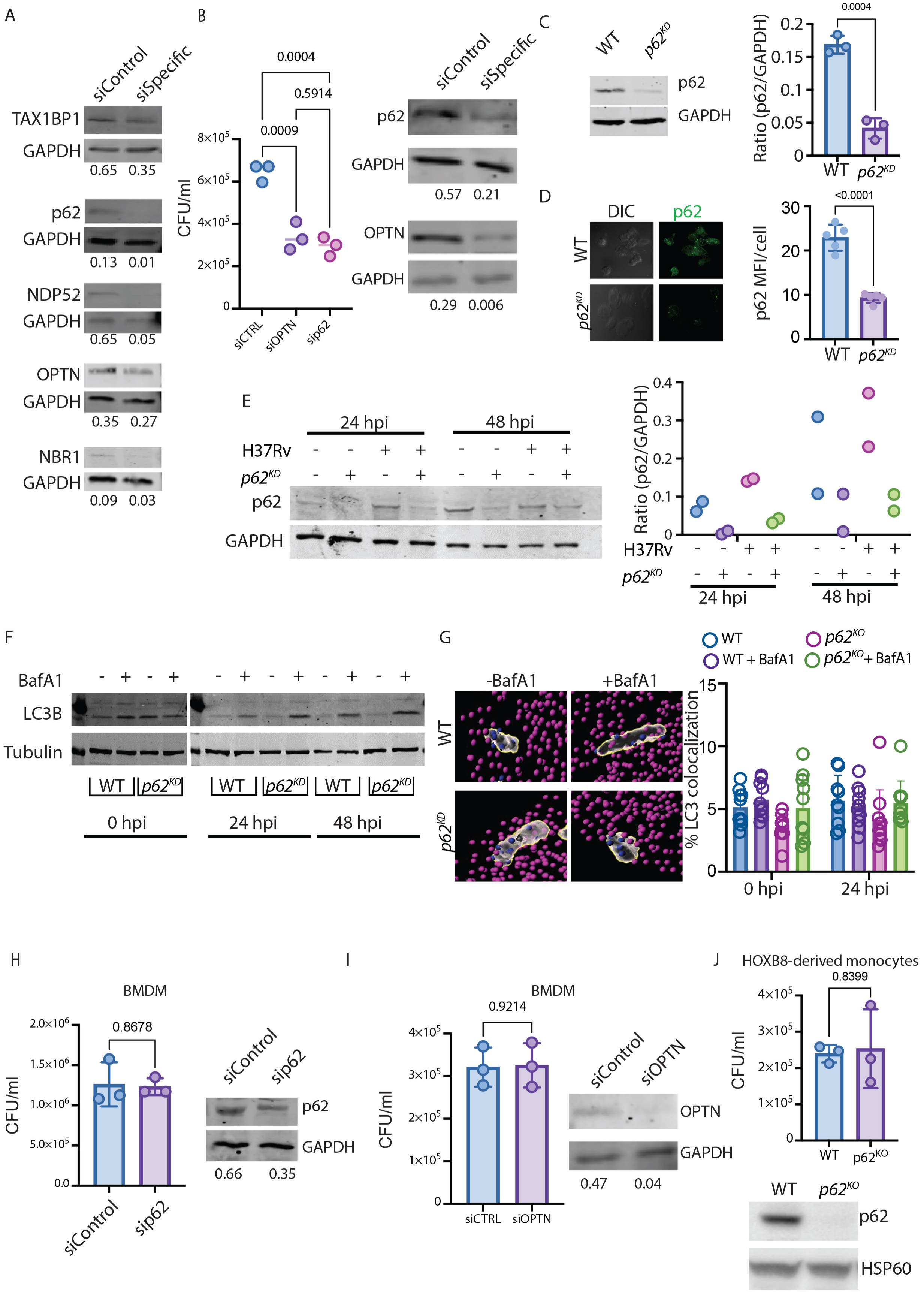
A
siControl
siSpecific
TAX1BP1
GAPDH
0.65 0.35
p62
GAPDH
0.13 0.01
NDP52
GAPDH
0.65 0.05
OPTN
GAPDH
0.35 0.27
NBR1
GAPDH
0.09 0.03
B
0.0004
0.5914
0.0009
CFU/ml
siCTRL
siOPTN
sip62
p62
GAPDH
0.57 0.21
OPTN
GAPDH
0.29 0.006
C
WT
p62KD
p62
GAPDH
Ratio (p62/GAPDH)
0.0004
D
DIC
p62
WT
p62KD
p62 MFI/cell
<0.0001
E
24 hpi
48 hpi
H37Rv
p62KD
p62
GAPDH
Ratio (p62/GAPDH)
F
BafA1
LC3B
Tubulin
WT
p62KD
0 hpi
24 hpi
48 hpi
G
WT
p62KO
WT + BafA1
p62KO+ BafA1
-BafA1
+BafA1
% LC3 colocalization
0 hpi
24 hpi
H
BMDM
0.8678
CFU/ml
siControl
sip62
p62
GAPDH
0.66 0.35
I
BMDM
0.9214
CFU/ml
siCTRL
siOPTN
siControl
siOPTN
OPTN
GAPDH
0.47 0.04
J
HOXB8-derived monocytes
0.8399
CFU/ml
WT
p62KO
p62
HSP60

**Fig. S2. Depletion of autophagy adaptors in human and mouse macrophages and their effect on H37Rv survival.**

**(A)** Immunoblot represent the siRNA-targeted depletion of adaptors TAX1BP1, p62, NDP52, OPTN and NBR1 in THP-1 monocytes in the separate experiment for each adaptor at 24 hours post-transfection. The numbers depict the ratio of intensities of specific proteins to that of GAPDH. **(B)** CFU analysis of H37Rv in control and siRNA-targeted-OPTN and -p62 U937 macrophages at 24 hpi. Right panel: Immunoblot represents the siRNA-targeted depletion of p62 and OPTN at 24 hours post-transfection. The numbers depict the ratio of intensities of specific proteins to that of GAPDH. **(C-D)** Respective immunoblot and IFA images for p62 in $p62^{KD}$ THP-1 cells generated using CRISPR/Cas9 and their quantification graphs (Right Panel). Plots represent data from three experiments (C) and 5 fields from two different experiments (D). **(E)** Immunoblot depicts the levels of p62 in uninfected and infected cells in WT and $p62^{KD}$ cells at 24- and 48 hpi and separate quantification plots from two independent experiments (Right). **(F)** Immunoblot represents LC3B levels in the lysates of WT and $p62^{KD}$ macrophages at 0-,24-, and 48-hpi in the presence and absence of BafA1 (100nM for 3 hours before processing). **(G)** Confocal images show the interaction of LC3B spots (blue spots, at <0.2 um from bacteria) and H37Rv. The bar graph represents the quantification of blue spots on H37Rv in WT and $p62^{KD}$ cells in the presence and absence of BafA1 at 0- and 24-hpi. Data show mean ± SD, n> 9 Fields, 20-35 per field, from three experiments. **(H-I)** Respective CFU analysis of H37Rv in control and siRNA-targeted p62/sqstm1 depleted and Optn depleted mouse bone marrow monocyte-derived macrophages at 24 hpi. Data show mean ± SD, from three independent experiments, right panel shows immunoblot confirming the p62 and OPTN depletion upon siRNA transfection. The numbers depict the ratio of intensities of specific proteins to that of GAPDH. **(J)** CFU analysis of H37Rv in WT and $p62^{KO}$ HOXB8 derived macrophages at 24 hpi. Data show mean ± SD, from three independent experiments. The immunoblot shows the levels of p62 in these cells.

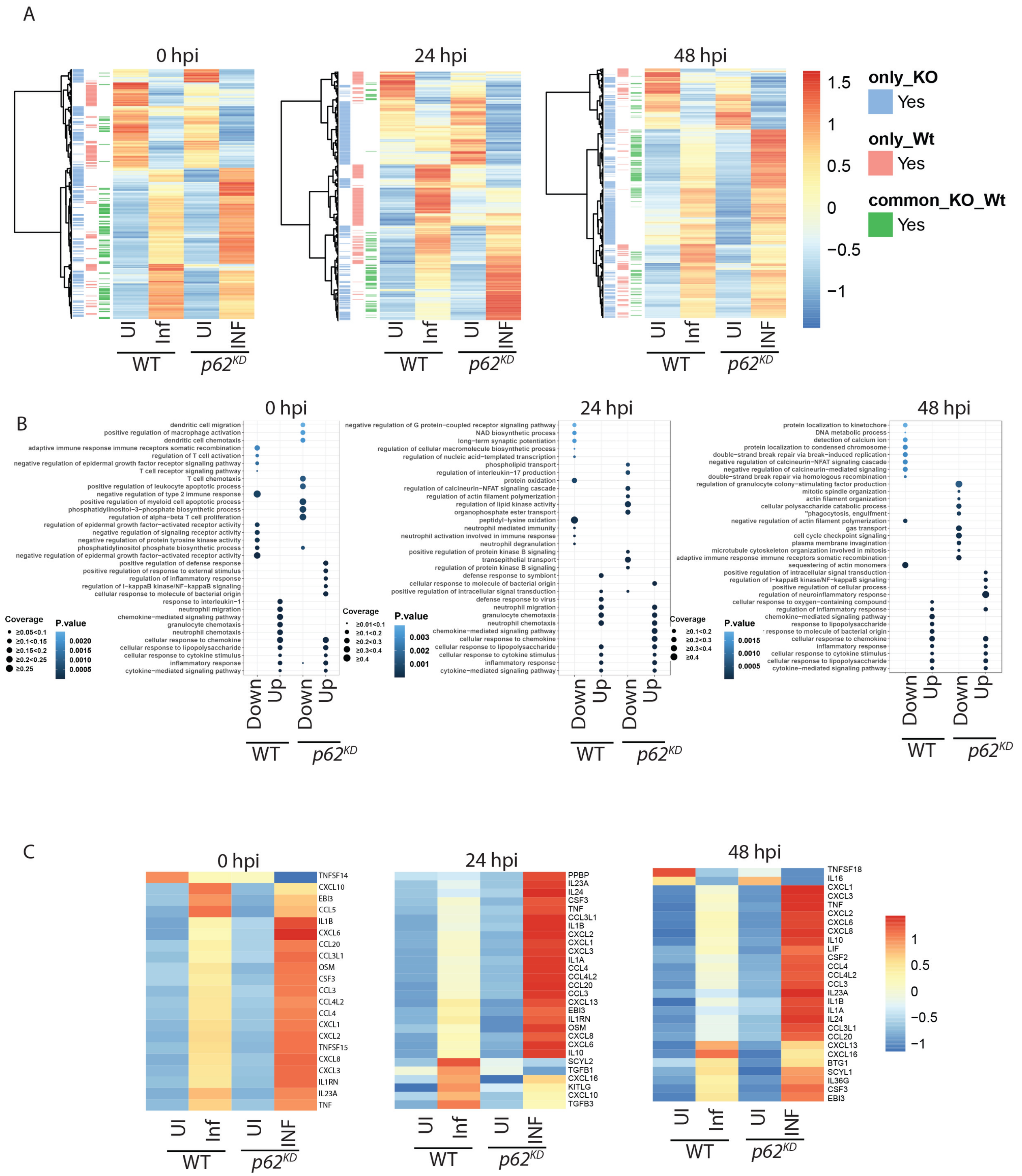


**Fig. S3.**

**Gene expression patterns and functional class enrichment. (A)** DEGs (differentially expressed genes, absolute log2FC ≥1 & q-value ≤ 0.1) at each time point. Row annotation represents DEGs identified when compared between infected versus uninfected in WT and *p62*$^{KD}$ macrophages at 0-, 24- and 48-.hpi. Row clustering was done using ward. D2 methodology on Euclidean distance calculated from the z-scaled values. Heatmap key reflects expression intensity based on z-scaled values **(B)** shows Gene Ontology Biological Process (GOBP) enrichment of the upregulated or downregulated genes identified based on infected versus uninfected in WT and $p62^{KD}$ macrophages at 0-, 24- and 48-.hpi. Significance is determined by nominal p.value ≤ 0.05. Only the top 15 enriched GOBPs from each comparison are visualised. Coverage represents the proportion of genes identified in a particular GOBP out of the total genes associated with that process. Colour intensity represents the nominal p-value gradient. **(C)** is a subset of **(A)** which shows an expression of the identified DE cytokine genes between infected versus uninfected in WT and $p62^{KD}$ macrophages at 0-, 24- and 48-.hpi

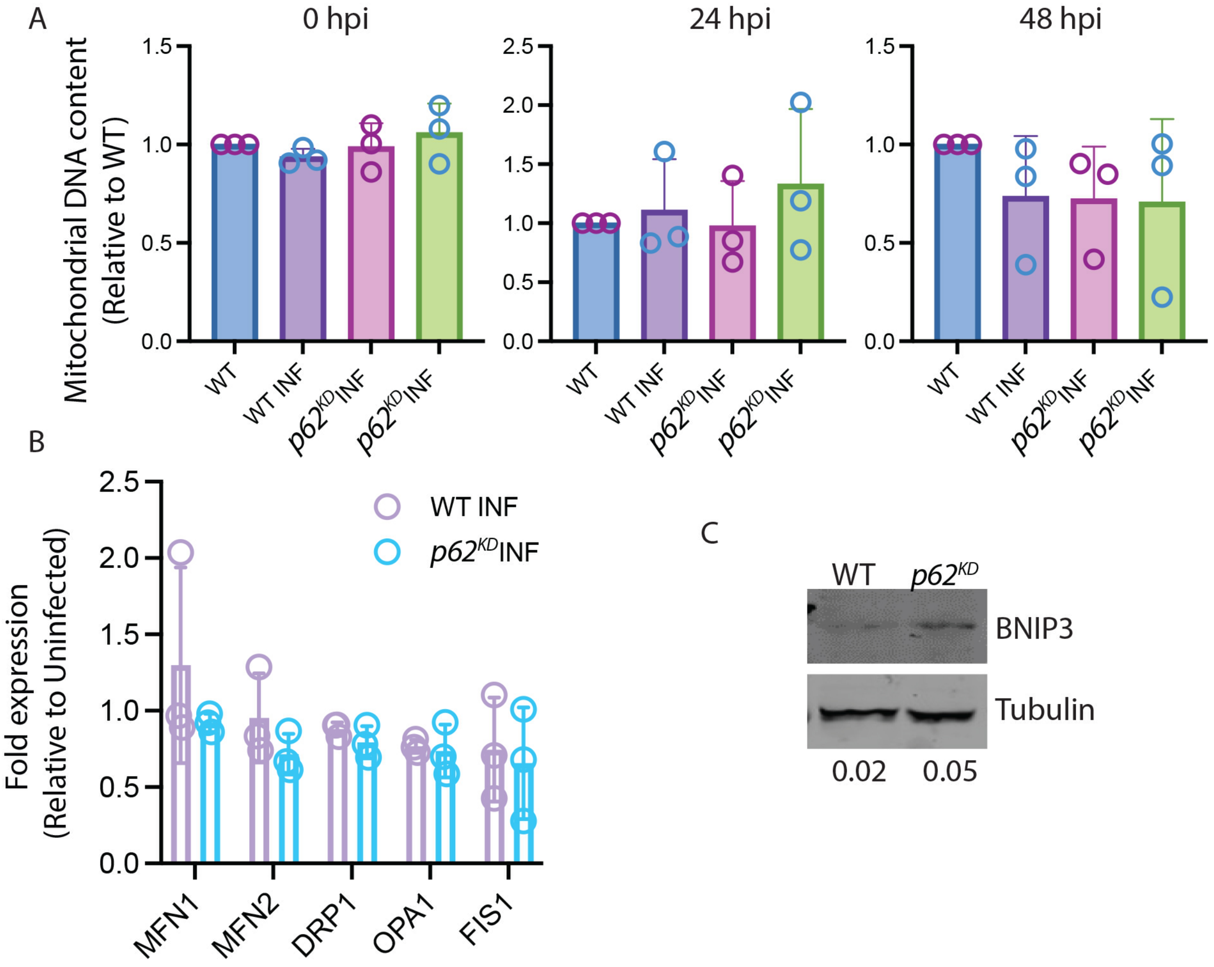


**Fig. S4.**

**Impact of p62/SQSTM1 depletion on mitochondrial DNA content and fission-fusion pathways (A)** Quantification of mtDNA by normalising the gene expression of the mitochondria-encoded protein (mt*CO1*) with nuclear genome encoded RNA polymerase (*RPL13A*) in WT and *p62*$^{KD}$ THP-1 macrophages in infected and uninfected THP-1 macrophages at 0-, 24-, and 48-hpi. **(B)** Fold change in gene expression of mitochondrial fusion proteins MFN1, MFN2 and OPA1, and fission proteins DRP1 and FIS1 by RT-PCR in H37Rv infected WT and *p62*$^{KD}$ THP-1 macrophages at 24 hpi. **(C)** Immunoblots represent protein expression of BNIP3 in infected WT and p62KD cells at 24 hpi.

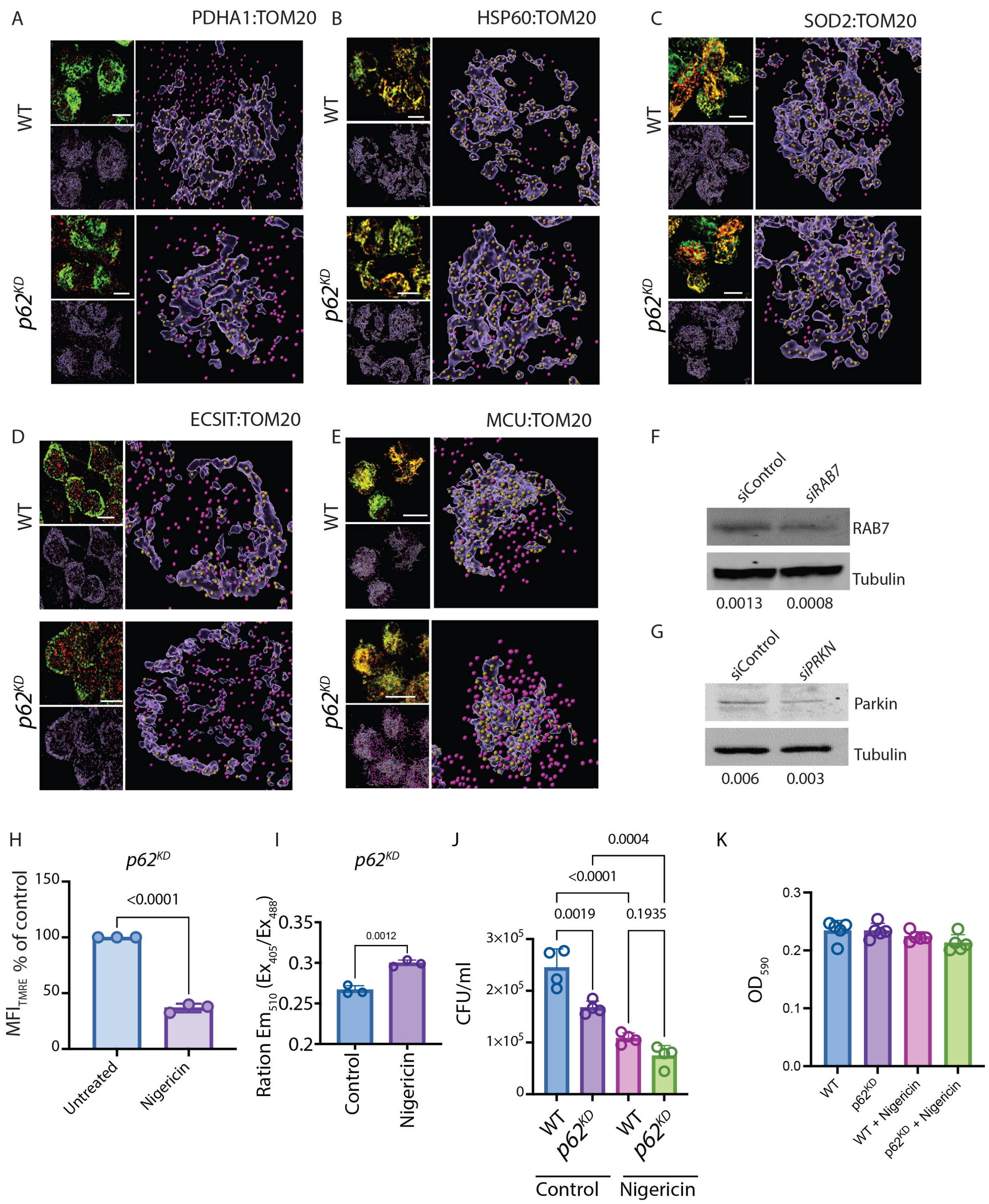


**Fig. S5.**

**Effect of mitochondrial quality control in H37Rv infected THP-1 macrophages (A-E)** Confocal images depict the interaction of TOM20 (green) with proteins from three different mitochondrial compartments (red). Representative TOM20 3D structures and mitochondrial protein spots: (A) PDHA1, (B) HSP60, (C) SO2, (D) ECSIT, and (E) MCU are shown at the right. The yellow spots are at <0.2 µm, and the magenta spots are at > 0.2 µm from TOM20 structures. **(F)** Immunoblot show the levels of RAB7 in siControl and siRAB7 in THP-1 macrophages at 24 hours post-siRNA transfection. **(G)** Immunoblot show the levels of PARKIN in siControl and si*PRKN* in THP-1 macrophages at 24 hours post-siRNA transfection. **(H)** Per cent MFI change in TMRE of infected THP-1 macrophages in the presence and absence of nigericin. **(I)** The relative oxidation state of *Mrx1-roGFP2* reporter as the ratios of emission MFI at 510nm when excited at 405nm to 488nm in Nigericin-treated WT and $p62^{KD}$ cells at 24 hpi. Data show mean ± SD, n=3. **(J)** H37Rv CFU in nigericin-treated and untreated WT and $p62^{KD}$ cells. Data show mean ± SD, n=4, from two independent experiments. (K) The graph represents the MTT cell viability assay of nigericin-treated and untreated WT and $p62^{KD}$ THP-1 macrophages.Scale bar: 10 µm.

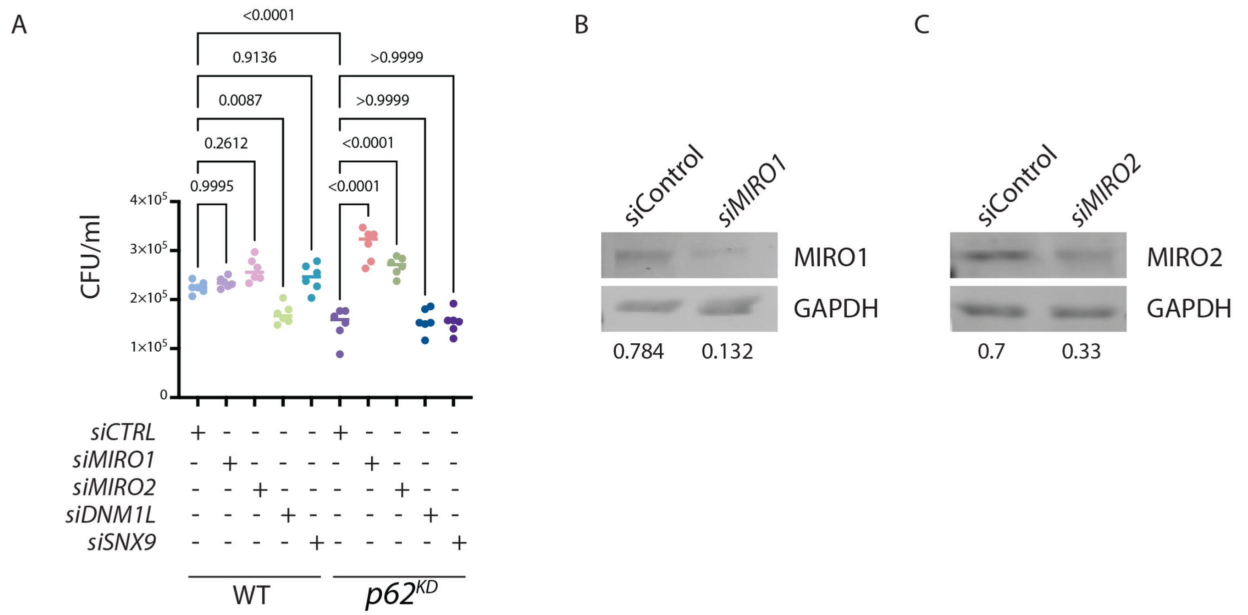


**Fig. S6.**

**Role of MIRO1 and MIRO2 in MDV biogenesis**

(A) CFU analysis of H37Rv in control and siRNA-targeted-MIRO1, MIRO2, DNM1L (DRP1) and SNX9 WT and *p62*$^{KD}$ macrophages at 24 hours post-transfection, Data show mean ± SD, n=5, from two independent experiments. (B-C) Immunoblot represents the siRNA-targeted depletion of MIRO1 and MIRO2, respectively at 24 hours post-transfection.

**Table S1.**
Infection-associated changes at different hours

**Table S2.**
Enrichment analysis of the DEGs

**Table S3.**
DEGs identified to be associated with mitochondria based on Mitocarta 3.0 (a subset of total DEGs)

**Table S4**:
Mitochondrial DEGs associated with mitochondria pathways based on Mitocarta 3.0

**Table S5:** DEGs identified to cytokine genes (a subset of total DEGs)
